# Dopamine-independent state inference mediates expert reward guided decision making

**DOI:** 10.1101/2021.06.25.449995

**Authors:** Marta Blanco-Pozo, Thomas Akam, Mark E. Walton

## Abstract

Rewards are thought to influence future choices through dopaminergic reward prediction errors (RPEs) updating stored value estimates. However, accumulating evidence suggests that inference about hidden states of the environment may underlie much adaptive behaviour, and it is unclear how these two accounts of reward-guided decision-making should be integrated. Using a two-step task for mice, we show that dopamine reports RPEs using value information inferred from task structure knowledge, alongside information about recent reward rate and movement. Nonetheless, although rewards strongly influenced choices and dopamine, neither activating nor inhibiting dopamine neurons at trial outcome affected future choice. These data were recapitulated by a neural network model in which frontal cortex learned to track hidden task states by predicting observations, while basal ganglia learned corresponding values and actions via dopaminergic RPEs. Together, this two-process account reconciles how dopamine-independent state inference and dopamine-mediated reinforcement learning interact on different timescales to determine reward-guided choices.

## Introduction

Adaptive behaviour requires learning which actions lead to desired outcomes and updating these preferences when the world changes. Reinforcement learning (RL) has provided an influential account of how this works in the brain, with reward prediction errors (RPEs) updating estimates of the values of states and/or actions, in turn driving choices. In support of this framework, dopamine activity resembles RPEs in many behaviours^1–4^, and causal manipulations can reinforce or suppress behaviours consistent with dopamine acting functionally as an RPE^5–9^.

However, value learning is not the only way we adapt to changes in the environment. For example, we behave differently on weekdays and weekends, but this is clearly not because we relearn the value of going to work versus spending time with family each Saturday morning. Rather, although the world looks the same when we wake up, we understand that the *state* of the world is in fact different, and this calls for different behaviour. Formally, the decision problem we face is partially observable – our current sensory observations only partially constrain the true state of the world. In such environments, it is typically possible to estimate the current state better using the history of observations than using just the current sensory input^10,11^ – we know today is Saturday because yesterday was Friday.

It is increasing clear that this ability to infer hidden (i.e., not directly observable) states of the world plays an important role even in simple laboratory reward-guided decision making^11–24^. For example, in probabilistic reversal learning tasks where reward probabilities of two options are anti-correlated, both behaviour and brain activity indicate that subjects understand this statistical relationship^11,17,18,25^. This is not predicted by standard RL models in which RPEs update the value of preceding actions, but is by models which assume subjects understand there is a hidden state which controls both reward probabilities. Intriguingly, brain recordings have shown that not only prefrontal cortex (PFC) but also the dopamine system can reflect knowledge of such hidden states^14,18,25–30^.

Integrating these two accounts of behavioural flexibility raises several pressing questions. If state inference, not RL, mediates flexible reward-guided behaviour, why does dopamine look and act like an RPE? Conversely, if value updates driven by dopaminergic RPEs are responsible, how does this generate the signatures of hidden state inference seen in the data?

To address these questions, we measured and manipulated dopamine activity in mice performing a two-step decision task, focusing on what we denote as ‘expert’ behaviour after task acquisition when mice were highly proficient at responding to changing reward contingencies. The task had two important features. First, reward probabilities were anti-correlated and reversed periodically, constituting a hidden state that could be inferred by observing where rewards were obtained. Second, inference and RL-based strategies could be differentiated by measuring how prior rewards affected dopamine activity. Behaviour was well explained by a model which inferred the hidden state of the task’s reward probabilities. Dopamine responses were consistent with RPE signalling, but critically the RPEs used value estimates informed by the inferred hidden state of the reward probabilities. Strikingly, though optogenetic stimulation of dopamine neurons was sufficient to reinforce immediately preceding choices, neither activating nor inhibiting dopamine neurons at trial outcome – the time when natural rewards elicited the strongest dopamine response – had any effect on subsequent choice. These apparently paradoxical data could be reproduced by a neural network model in which a recurrent PFC network learned to track the hidden state of the reward probabilities by predicting observations, and a feed-forward basal-ganglia network learned appropriate actions based on this inferred state using RL.

This intimates that RL and inference-based accounts of reward-guided decision making can be reconciled in a combined picture where the effect of rewards on choices in well-learned tasks are mediated by changes in recurrent PFC activity representing hidden states, not RPEs, but dopaminergic RPEs across task acquisition are critical for learning the values and actions these states imply.

## Results

### Mice behaviour respects task causal structure

We trained DAT-cre mice (n=18) on a sequential decision task, which required them to choose between two options – left and right – to gain access to one of two reward ports – top or bottom (Figure 1A). Each trial started with a central port lighting up, which mice poked to initiate the trial. This triggered either a choice state, where both left and right ports lit up (75% of trials), or a forced choice where either left or right lit up (25% of trials). Poking an illuminated side port triggered a transition to one of two possible second-step states, signalled by a 1-s auditory cue, in which either the top or bottom port was illuminated. Each first-step choice (left or right) led commonly (80% of trials) to one second-step state (up or down) and rarely (20% of trials) to the other (Figure 1B). Poking the active reward port caused a 500-ms auditory cue indicating the trial outcome (reward or not), with reward delivered at cue offset. Reward delivery in each port was probabilistic, with probabilities that changed in blocks between *0.8/0.2, 0.5/0.5* and *0.2/0.8* on the up/down ports respectively (Figure 1C bottom). The reward probabilities were therefore anticorrelated at the up and down ports. Though we have previously used a two-step task for mice where both transition and reward probabilities changed over time^31,32^, here the transition probabilities were fixed and only reward probabilities changed. This facilitates hidden-state inference strategies (also referred as ‘latent-state’ inference^33^) because the state of the reward probabilities – whether up or down is rewarded with higher probability – fully determines which first-step action is best^31,33^.

**Figure 1.**
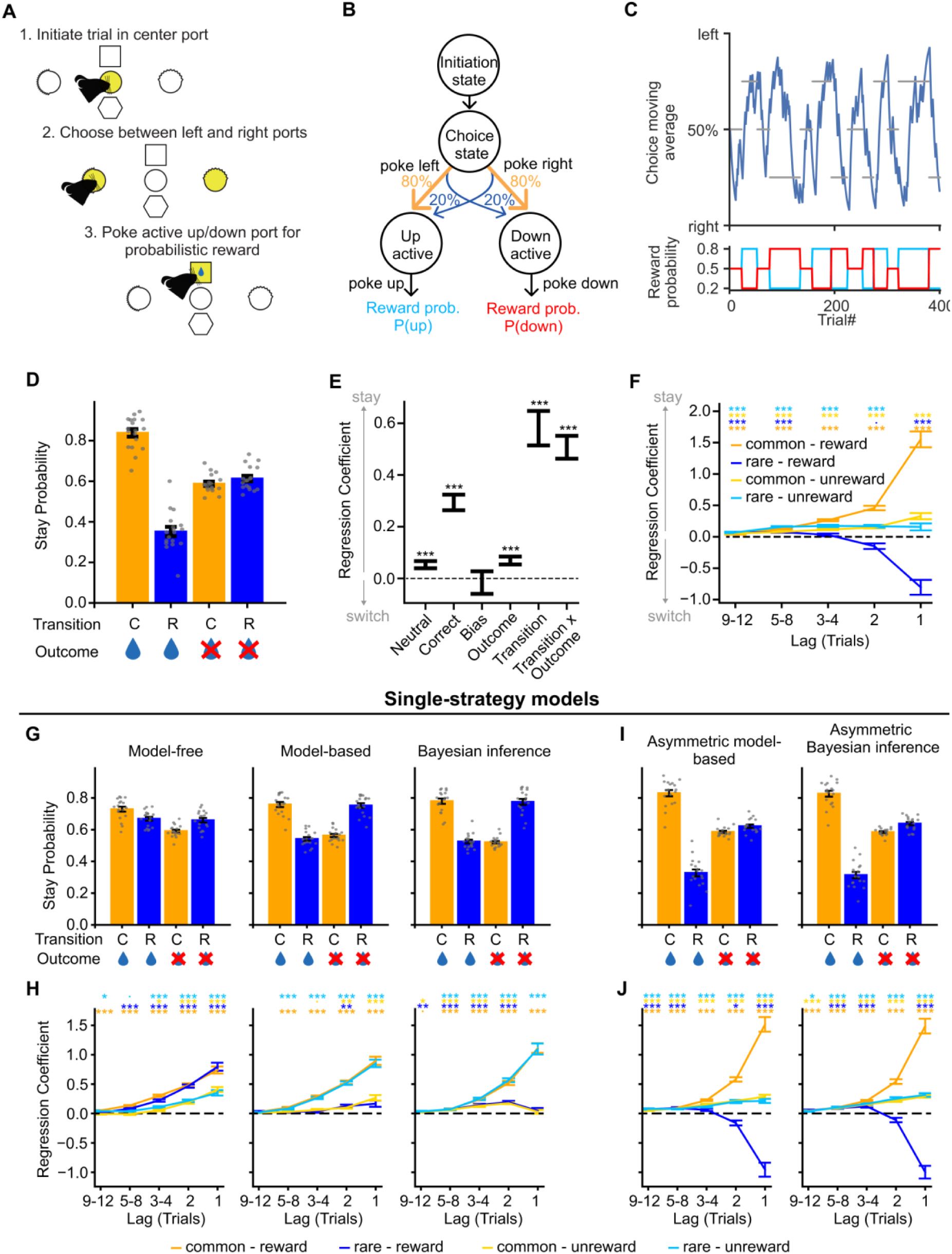
Two step task behaviour. **A)** Diagram of trial events. **B)** Diagram of the task state space, reward and transition probabilities. **C)** Example behavioural session. Top panel: Moving average of choices (blue trace) with reward probability blocks shown by grey bars, indicating by their vertical position whether the reward probability was higher for the state commonly reached for the left or right choice or a neutral block. Bottom panel: Reward probability at the up (blue) and down (red) ports. **D)** Probability of a choice being repeated as a function of the subsequent state transition (common or rare) and trial outcome (rewarded or unrewarded). Dots show individual subjects, error bars cross subject SEM. **E)** Mixed effects logistic regression predicting repeating choice as a function of the trial outcome (rewarded or not), state transition (common or rare), the transition-outcome interaction, and whether the choice was to the correct (high reward probability) option. **F)** Lagged logistic predicting choice as a function of the history of different trial types defined by the transition and outcome. **G-J)** Single strategy models: stay probability (as in Figure 1D) (**G, I**) and lagged regression (as in Figure 1F) (**H, J**) of simulated behaviour from different RL models fitted to mice behaviour. *p<0.05, **p<0.01, ***p<0.001

Subjects tracked which option was currently best, performing 348.70 ± 93.90 (mean ± SD across subjects) trials and completing 8.32 ± 3.24 (mean ± SD across subjects) blocks per session (Figure 1C). Rewards did not simply reinforce the choice made at the start of the trial. Instead, reward following a common transition promoted repeating the same first-step choice on the next trial, while reward following a rare transition promoted switching to the other first-step choice (Figure 1D,E, mixed effects logistic regression – transition x outcome: β = 0.507, SE = 0.044, z = 11.468, p < 0.001), and these effects persisted over multiple trials (Figure 1F). This pattern is adaptive because it corresponds to rewards promoting choice of first-step actions that commonly lead to the second-step states where they were obtained. However, the probability of repeating the same choice following a non-rewarded outcome was similar irrespective of whether a common or rare transition occurred (Figure 1D,F). This strong asymmetry between the effect of reward and reward omission is surprising given that both outcomes are equally informative about future reward, but might reflect differences between task statistics and the demands of foraging in natural environments.

To assess what strategy the animals used, we fit a set of models to their choices and simulated data from the fitted models (Figure 1G-I). Neither model-free nor model-based RL, commonly used to model two-step task behaviour^34^, resembled subject’s choices (Figure 1G, H). We also considered a strategy that used Bayesian inference to track the hidden state of the reward probabilities, combined with a fixed habit-like mapping from this state estimate to the corresponding high-value first-step action^33^, but again this did not resemble the experimental data. The reason these models failed is that, though model-based RL and inference strategies can reproduce our experimental finding that rewards reinforced the first-step action that commonly leads to the state where the reward was obtained^33,34^, both predict a *symmetric* influence of reward and omission on choices, contrary to our experimental data.

We therefore modified each model to incorporate this asymmetry. For model-based RL this was done using different learning rates for positive and negative RPEs. We also incorporated forgetting about the value of states that were not visited, as this was supported by model comparison (Figure S1). This approach was not possible for the inference model, as Bayesian updates do not have a learning rate parameter that can be different for reward and omission. We therefore implemented the asymmetry by modifying the observations that the model received from the task. Specifically, the asymmetric inference model treated reward obtained in the two second-step states as different observations, but treated reward omission as the same observation irrespective of the state where it occurred. Simulated on the task, both asymmetric models generated a pattern of stay probabilities that closely matched subject’s data (Figure 1I, J).

We adopted two different approaches to try and differentiate between these strategies: i) likelihood-based model comparison, and ii) fitting a mixture-of-strategies model incorporating both components to assess which explained most variance in subjects’ choices. Both analyses gave a consistent picture that it was not possible to arbitrate between the strategies using behaviour alone (Figure S1). Critically, however, the two strategies make different predictions for how rewards update the estimated value of each second-step state (discussed below), and hence for dopaminergic RPE signalling. We therefore looked for evidence of inference-based value updates in dopamine activity.

#### Inferred values drive dopamine signals

We used fibre photometry to record calcium signals from GCaMP6f expressing dopamine neuron cell bodies in the ventral tegmental area (VTA) and axons in nucleus accumbens (NAc) and dorsomedial striatum (DMS) (Figure 2A), and dopamine release using dLight1.1 expressed pan-neuronally in NAc and DMS (Figure 2B, see Figure S2 for placements). Dopamine activity fluctuated dynamically across the trial, as mice made their initial choice, received information about the second-step state reached, and trial outcome (Figure 2D-F). Reward responses were prominent in all signals, though relatively weaker in DMS calcium. However, average DMS calcium activity masked a strong medio-lateral gradient in reward response, with larger responses more laterally in DMS (Figure S3). For the following analyses, we excluded the DMS site in 2 animals where the fibre was most medial and we observed a negative reward response (Figure S3).

**Figure 2.**
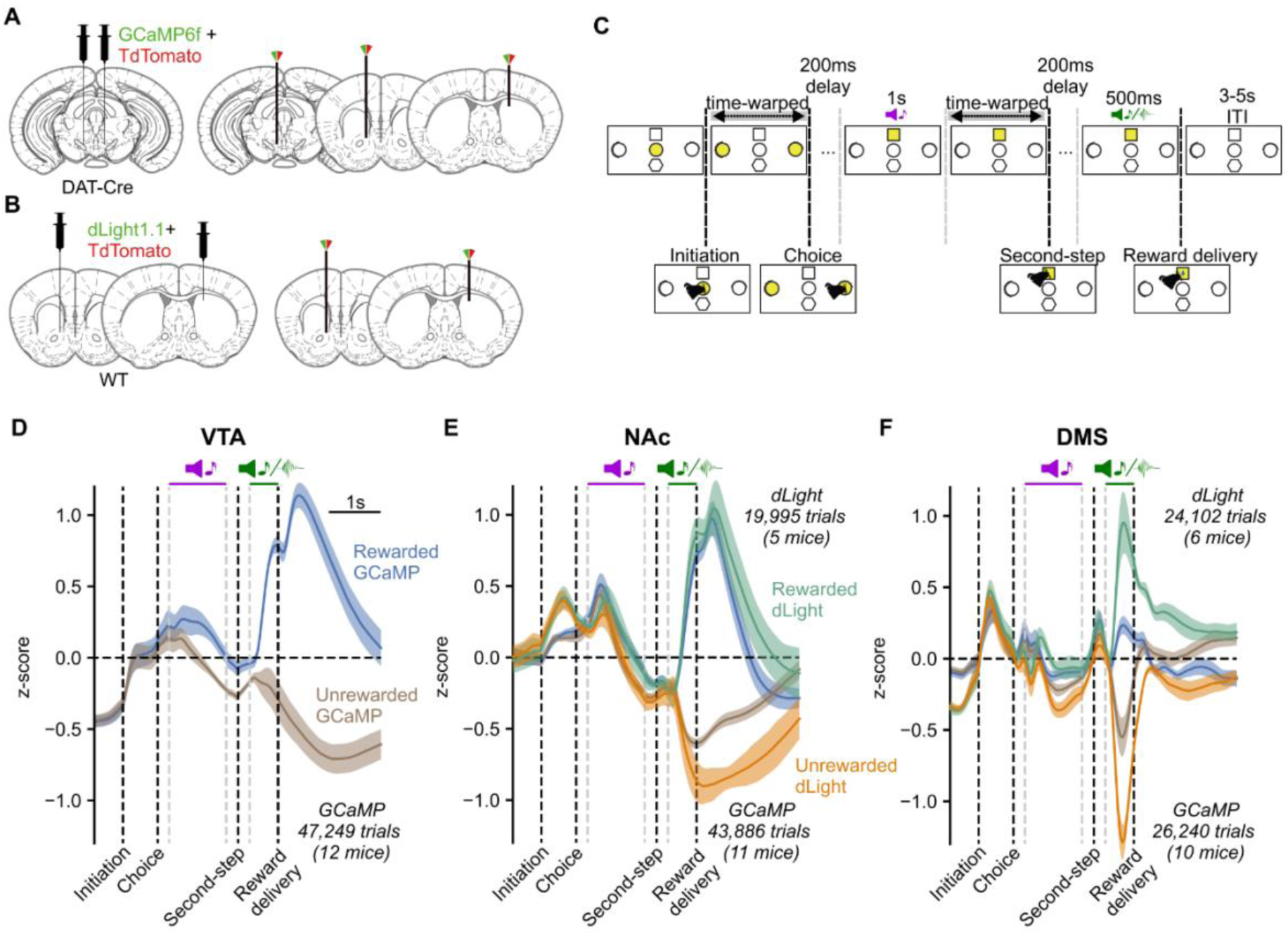
Dopamine activity and release are modulated across the trial. **A-B)** Injection and implant schematic. **A)** For calcium photometry, DAT-Cre mice were injected bilaterally with GCaMP6f and TdTomato in VTA, and optic fibres were implanted in VTA and NAc in one hemisphere, and DMS in the other hemisphere. **B)** For dLight photometry, WT mice were injected with dLight1.1 and TdTomato in NAc and DMS in different hemispheres, and optic fibres implanted at the same sites. **C)** Trial timeline. Trials were aligned by warping the intervals between the trial initiation and choice, and between the second-step port lighting up and being poked. **D-F)** Mean z-scored calcium signal and dopamine receptor binding on rewarded and unrewarded trials with shaded areas indicating cross-subject standard error in **D)** VTA GCaMP (n: 12 mice, 121 sessions, 47,249 trials), **E)** NAc GCaMP (n: 11 mice, 108 sessions, 43,886 trials) and dLight (n: 5 mice, 64 sessions, 19,995 trials), **F)** DMS GCaMP (n: 10 mice, 74 sessions, 26,240 trials) and dLight (n: 6 mice, 91 sessions, 24,102 trials).

A key feature of the state inference strategy is that it assumes that a single hidden variable controls both reward probabilities. Therefore, reward obtained in one second-step state not only *increases* the value of that state but also *decreases* the value of the other second-step state, unlike in standard model-based RL where the state values are learned independently (Figure 3C). We can therefore leverage our photometry data to discriminate between these strategies by examining how the previous trial’s outcome influences dopamine activity and release when the second-step state reached on the current trial is revealed. Specifically, we can ask whether a reward obtained in one second-step state (e.g., *up-active*)*, decreases* the dopamine response to the other second-step step state (*down-active)* if it is reached on the next trial.

**Figure 3.**
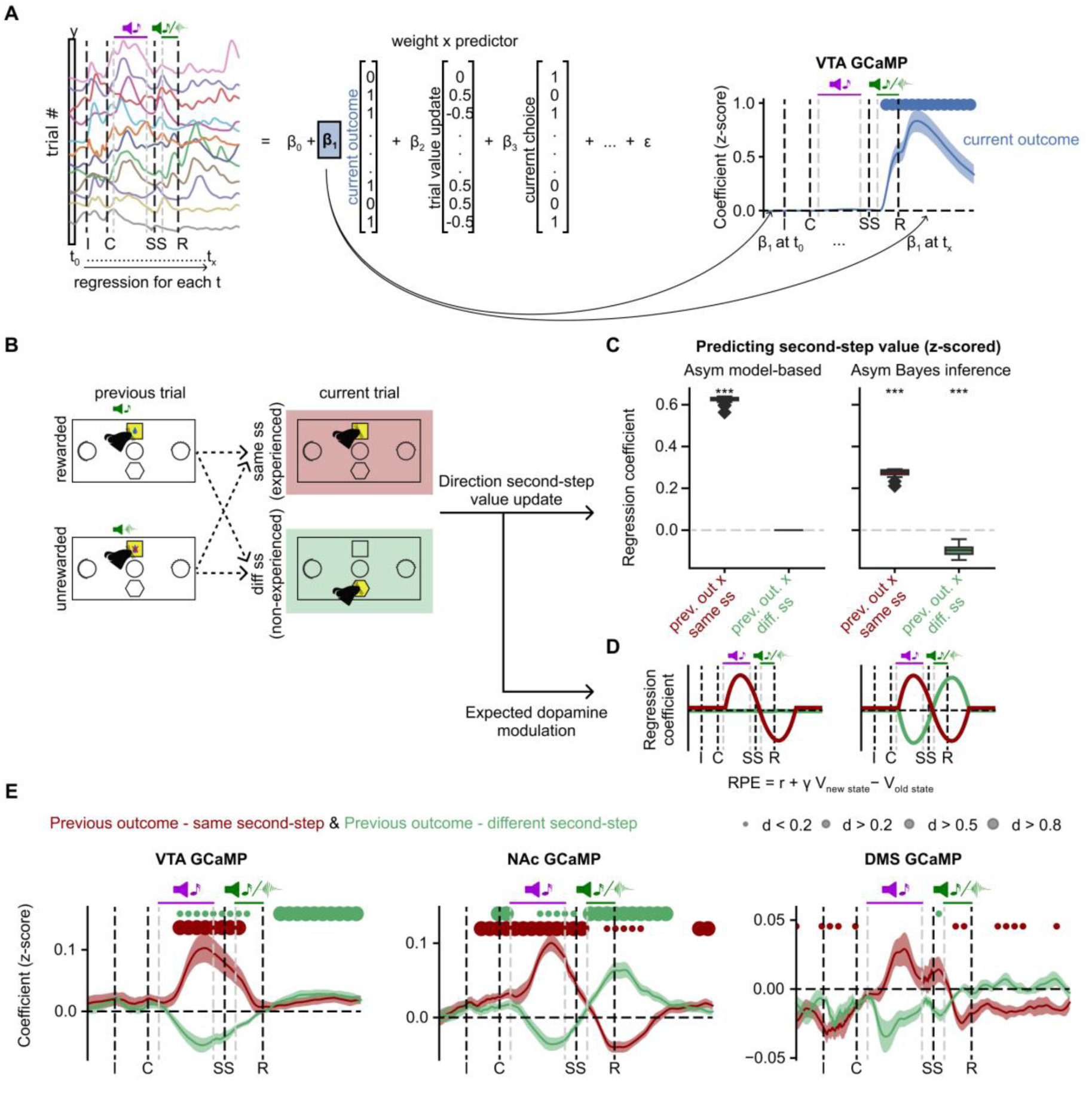
Inferred value information drives dopamine activity. **A)** Left, schematic of the linear regression model predicting dopamine activity for each timepoint in the trial. Right, coefficients in the linear regression for the current trial outcome predictor in VTA GCaMP signal, showing cross subject-mean and standard error. Dots indicate effect size at timepoints where coefficients are statistically significant, assessed by a two-sided t-test comparing the cross-subject distribution against 0, Benjamini-Hochberg corrected for comparison of multiple time-points. **B)** Schematic of the two-regressors used to test the influence of previous trial outcome on the value of the same (dark red) or different (green) second-step reached on the current trial. **C)** Linear regression predicting second-step state value as a function of the previous trial outcome, and whether the current trial second-step state was the same or different from the previous trial, for the model-based (Asym model-based) and Bayesian inference (Asym Bayes inference) strategies. The regression also included the other predictors used to explain photometry signal in A. **D)** Predicted dopamine modulation in the linear regression in A, based on the direction of the second-step value update from C under the assumption that dopamine modulation is consistent with the canonical RPE framework (where the value of the new state – i.e., second-state value at second-step cue – has a positive influence on dopamine activity while the value of the previous state – i.e., second-state value at outcome time – has a negative influence on dopamine activity). **E)** Coefficients in the linear regression predicting dopamine activity for the predictors in B, showing the influence of previous trial outcome when the second-step state was the same (dark red) or different (green) from the previous trial in VTA, NAc and DMS (See Figure S3,4 for dopamine concentrations, dLight). As in **A** right, showing cross subject-mean and standard error as the shaded area. Dots indicate effect size of the statistically significant timepoints, Benjamini-Hochberg corrected.

To do so, we aligned activity across trials and used linear regression to model dopamine fluctuations across trials and time-points. We ran separate regression analyses for each timepoint in the aligned data, using predictors which varied from trial-to-trial but took the same value for all timepoints in each trial. The time courses of predictor loadings across the trial, therefore, reflect when, and with what sign, each predictor explained variance in the activity (Figure 3A). The key predictors for differentiating between strategies are one coding for the previous trial’s outcome on trials where the second-step state is the *same* as on the previous trial, and another coding the previous trial’s outcome when the second step reached on the current trial is *different* (Figure 3B). We also included regressors modelling the current trial outcome and other possible sources of variance (see *Methods*, Figure S4, S5). We focus on the GCaMP data in the main figures, but results from dLight were closely comparable, except where noted in the text (see Figure S4, S5).

When the second-step was the <u>same</u> as on the previous trial, the previous trial’s outcome *positively* influenced dopamine when the second-step state was revealed, consistent with both model-based RL and inference (Figure 3D,E, S4). However, when the second-step state was <u>different</u> to that on the previous trial, the previous outcome *negatively* influenced dopamine when the second-step state was revealed. This is at odds with the predictions of standard model-based RL but, crucially, is consistent with inference (Figure 3D,E, S4). In NAc, loading on these regressors reversed sign at outcome time. This biphasic response is exactly as predicted for an RPE sign: RPE’s are computed from value differences between successive timesteps, so if dopamine reports RPE, the value of the second-step state reached on the trial will drive a positive response when the state is revealed, followed by a *negative* response at the time of trial outcome^35–37^. Unexpectedly, this reversal was not observed in either VTA or DMS.

To ensure that this pattern of modulation by inferred value was not an artefact of confounding effects of the trial history, we performed a lagged regression predicting the dopamine response to the second-step state cue. This confirmed that the dopamine response was driven by inferred state values, and that these integrated outcomes over multiple previous trials (less clearly in the DMS axonal calcium activity, but prominently in DMS dopamine release) (Figure S4, S6A,B).

To test whether the asymmetric influence of rewards and omissions on subject’s choices was also reflected in dopamine activity, we ran a modified regression analysis which used separate regressors for trials following rewarded and non-rewarded outcomes, each coding for whether the second-step state was the same or different as the previous trial (Figure S6C). In VTA and in NAc, the differential dopamine response to reaching same vs different second-step state was much stronger following rewarded than non-rewarded trials, consistent with rewards updating second-step state values more strongly than omissions (Figure S6C).

We were also able to resolve an influence of the previous outcome on dopamine activity at the time of first-step choice (Figure S4). Specifically, dopamine activity at choice time was higher when subjects choose an action that commonly led to the state where reward was obtained on the previous trial, consistent with subjects inferring the value of the first-step action using knowledge of the task structure. Again, these effects were primarily driven by rewarded rather than omission trials (Figure S6D).

Together, these findings indicate that the mice understood that a single hidden variable controlled the reward probabilities in both ports and inferred its current state by observing where rewards were obtained. Reward predictions based on this then shaped dopamine responses to both task states and subject’s choices.

#### Dissociable influence of RPE, reward rate and movement on dopamine activity

Recent work has argued that dopamine fluctuations more closely reflect value than RPE^38,39^. To examine whether this was the case in our data, we used the same linear regression framework but with value estimates from the inference model for the chosen first-step action and second-step state as predictors (Figure 4, S7, S8). As lateralised movements^6^ and average reward rate^5,38^ have also been reported to influence dopamine, we additionally included regressors for these variables.

**Figure 4.**
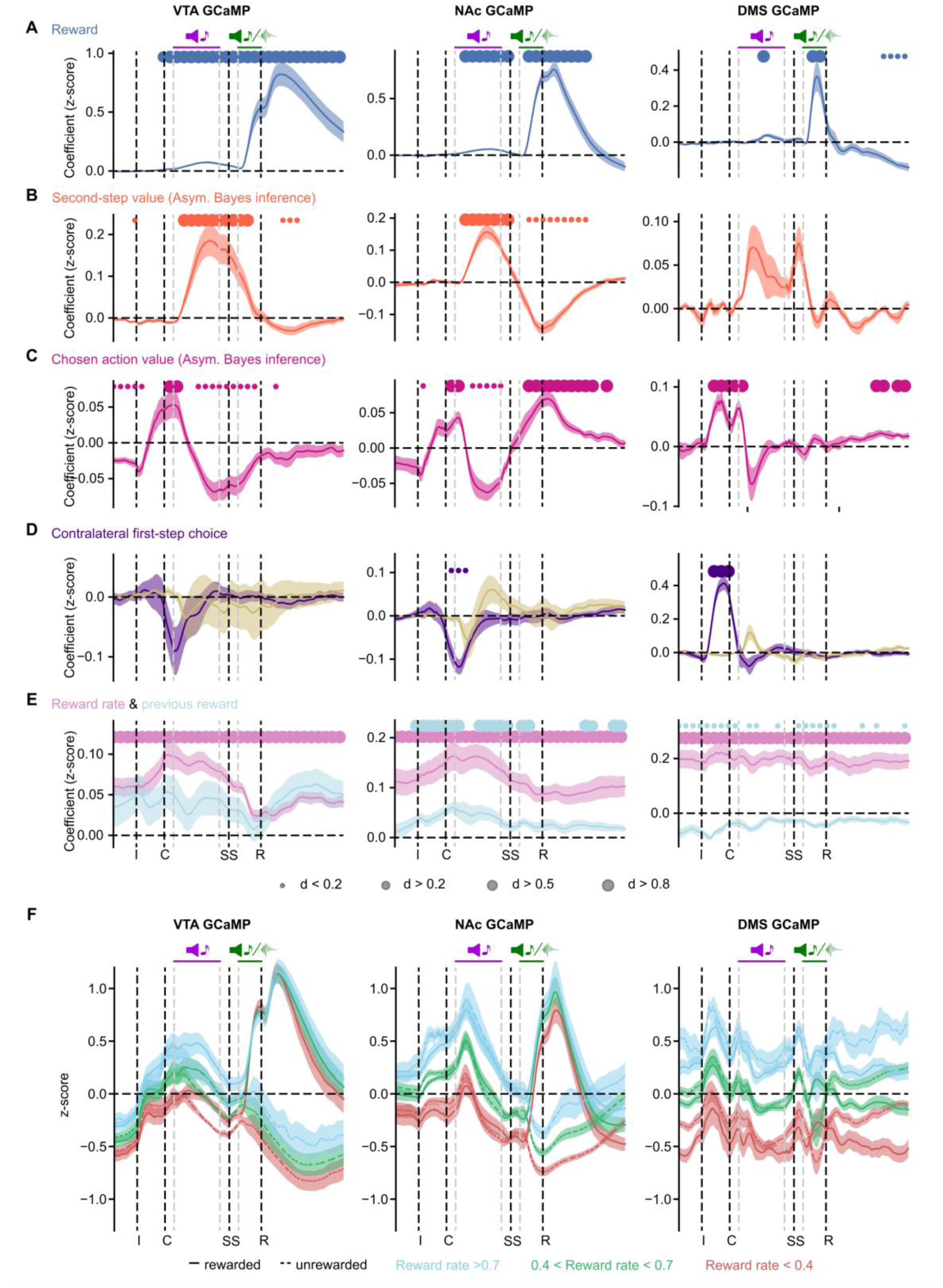
Simultaneous RPE, reward rate and lateralised movement signals in dopamine activity. **A-E)** Coefficients from the regression using the value estimates from the Bayesian inference model, predicting dopamine activity for each timepoint in the trial for each region, showing cross-subject-mean and standard error. Dots indicate effect size at timepoints where coefficients are statistically significant, assessed by a two-sided t-test comparing the cross-subject distribution against 0, Benjamini-Hochberg corrected for comparison of multiple time-points. **A)** Reward, Current trial outcome. **B)** Value of the second-step reached on the current trial derived from the asymmetric Bayesian inference model. **C)** Value of the chosed first-step action derived from the asymmetric Bayesian inference model. **D)** Contralateral first-step choice (coded according to the initial choice – left/right – relative to the hemisphere being recorded from). **E)** Recent reward rate (exponential moving average with tau = 8 trials), and previous trial outcome. **F)** Mean-z-score activity split by rewarded/unrewarded trials and recent reward rate. Blue – high reward rate > 0.7 rewards/trial (exponential moving average with tau = 10 trials). Green, medium reward rate (between 0.4 and 0.7). Red, low reward rate (less than 0.4).

In line with our above results, second-step state values drove a biphasic response in NAc GCaMP and dLight signals, with a positive influence when the second-step state was revealed followed by a negative influence at outcome time, consistent with RPE not direct value coding (Figure 4B). VTA GCaMP also showed this biphasic pattern but with a smaller negative response at outcome time relative to the positive response to the second-step state. The time-course was more complex in DMS, with peaks following both the second-step cue and second-step port entry, though the former only survived multiple comparison correction across timepoints in the dLight data (Figure S8). Chosen action value also had a strong positive influence on activity in all 3 regions around the time of the choice, which then reversed in all the regions when the second-step was revealed, again consistent with RPE (Figure 4C).

In addition to these RPE-like signals, dopamine was also transiently modulated by lateralised movement (Figure 4D, S4, S5, S7, S8). Consistent with previous reports^6,40^, activity in DMS but not VTA or NAc was modulated after initiating a contralateral choice. Unlike previous studies, here the task necessitated a second lateralised movement (in the opposite direction) from choice port back to the centrally located second-step reward port. Unexpectedly, this did not evoke a response in DMS, but did in VTA and NAc activity (note, the negative predictor loadings for VTA and NAC in Figure 4D following the choice indicates *increased* activity for contralateral movements from the choice port back to the second-step port).

Reward rate, calculated as an exponential moving average with time constant of 8 trials had a strong positive influence on dopamine in all 3 regions (Figure 4E, S4, S5, S7, S8) (similar results were obtained using time constants of 5-15 trials, data not shown). Unlike the influence of action/state values and rewards, which were tightly time-locked to the corresponding trial events (Figure 4A-C), reward rate positively influenced activity at all timepoints, with little modulation by specific trial events (Figure 4E, F). This reward rate signal was also present in NAc dopamine concentrations, but negligible in DMS concentrations (Figure S8).

In sum, these data demonstrate that dopamine carries information about i) action and state values in a manner consistent with RPE signalling, ii) lateralised movement, and (iii) recent reward rate. While these signals exist in parallel, they can nonetheless be dissociated based on their timescale and their lateralisation.

#### Dopamine does not mediate the reinforcing effect of task rewards

The above data demonstrate that dopamine reported RPE and reward information, but cannot establish whether this signal was causally responsible for updating mice’s subsequent choice behaviour. To assess this, we therefore manipulated dopamine activity in a new cohort of mice expressing either channelrhodopsin (ChR2, N=7) or a control fluorophore (EYFP, N=5) (Figure 5A, S9A) in VTA dopamine neurons. We verified that our stimulation parameters (5 pulses, 25 Hz, ∼8-10 mW power) were sufficient to promote and maintain intracranial self-stimulation in the ChR2 group compared to YFP controls (t(10) = 3.107, p = 0.011, 95% CI [68.18, 414.07], Cohen’s D = 1.819) using an assay where optical stimulation was delivered contingent on nose-poking in a different context from the two-step task (Figure 5B).

**Figure 5.**
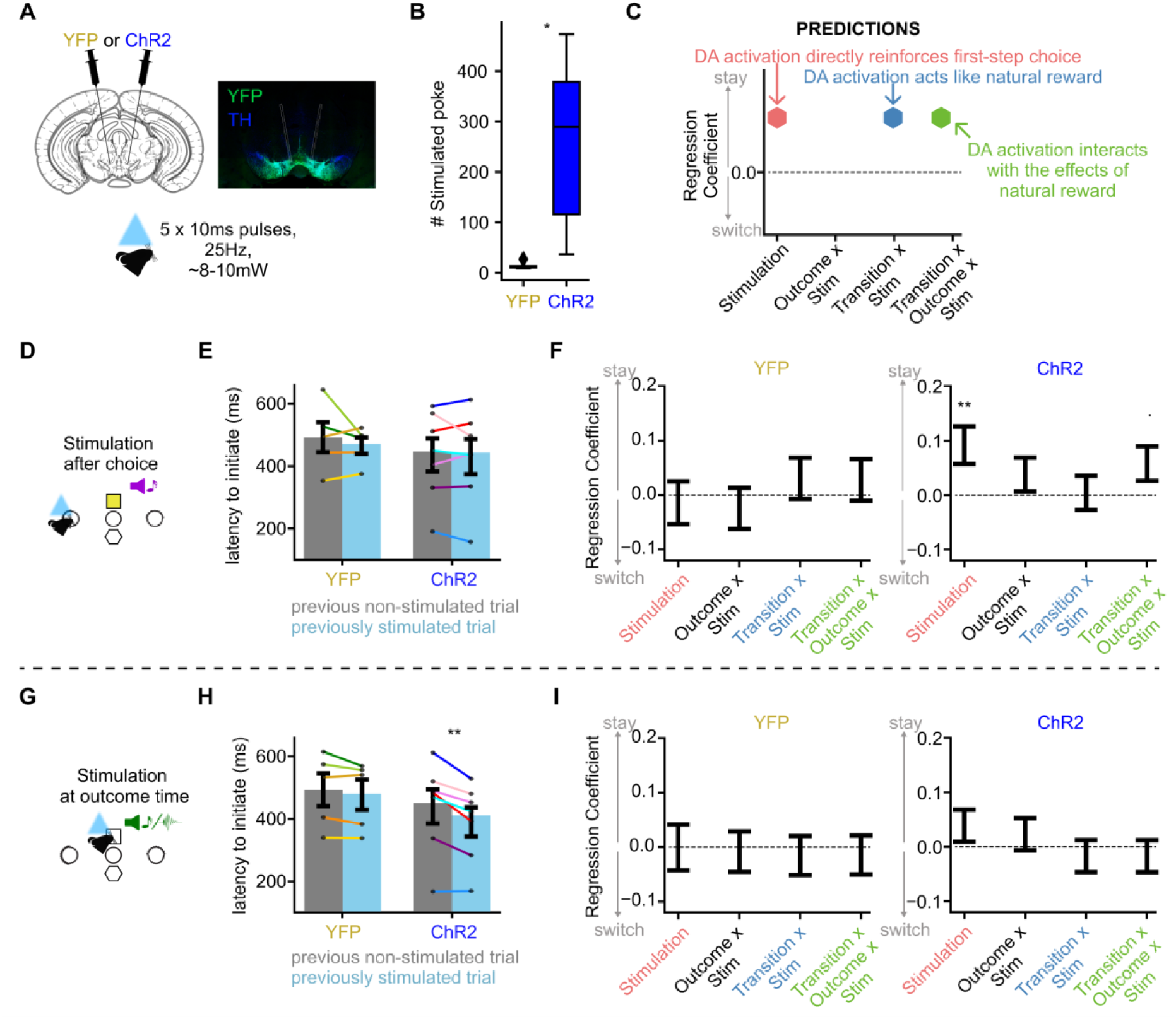
Dopamine stimulation does not recapitulate natural reward effects. **A)** chematic (left) and photomicrograph (right) showing injection and fibre placement. Photomicrograph comes from an example ChR2 mouse, stained for TH (Tyrosine Hydroxylase, blue) and YFP (yellow fluorescent protein). Stimulation consisted of 5 pulses at 25Hz, ∼8-10mW power. **B)** Intracranial self-stimulation. Mean number of entries to a nose-poke port delivering stimulation in the YFP vs the ChR2 group. Error bars show cross subject standard error. *p<0.05 two-sided independent samples t-test. **C)** Predictions on the effect of VTA dopamine optogenetic activation on animals’ subsequent choices. **D-F)** Effect of optogenetic stimulation 200ms after choice in the control (YFP) and ChR2 group. **G-I)** Effect of optogenetic stimulation at time of the trial outcome cue. **D,G)** Optogenetic stimulation schematic. **E-H)** Mean latency to initiate a new trial after the centre poke illuminates following stimulated and non-stimulated trials. Dots indicate individual animals, error bars show standard error. **p<0.01, two-sided paired t-test. **F,I)** Mixed effects logistic regression predicting repeating the same choice on the next trial using the regression model from Figure 1E, with additional regressors modelling the effect of stimulation and its interaction with trial events. Only regressors showing stimulation effects are shown here. See Figure S9 for the full model including both groups (YFP and ChR2) and the three stimulation types (non-stim, stim at choice, stim at outcome. ^•^p<0.1, **p<0.01. N: YFP: 5 animals, 13,079 trials (choice-time stimulation sessions), 15,419 trials (outcome-time stimulation sessions); ChR2: 7 animals, 20,817 trials (choice-time stimulation sessions), 23,051 trials (outcome-time stimulation sessions).

We then examined the effect on two-step task behaviour of optogenetic stimulation at two different timepoints – (i) after the first-step choice, at the time of second-step cue onset, or (ii) at outcome cue presentation (Figure 5D,G) (25% stimulated trials, stimulation timepoint fixed for each session, counterbalanced across sessions and mice, N=6-8 sessions per mouse and stimulation type). We again used a mixed effects logistic regression approach (Figure 1E), here adding stimulation and its interaction with transition and outcome as regressors. Note that if the effects of rewards are mediated by dopamine, we would expect stimulation to act like a reward, causing positive loading on the *transition x stimulation* regressor, or, if stimulation interacts with outcome (reward/omission), on the *transition x outcome x stimulation* regressor; if, however, optogenetic activation simply reinforces the previous choice, this would be evident as positive loading on the *stimulation* regressor (Figure 5C).

Data from both groups (YFP and ChR2) and each stimulation type (non-stimulated trials, stimulation after first-step choice, and stimulation at outcome) were included in a mixed-effects logistic regression. This revealed a significant stimulation-type1-by-group effect (β = -0.053, SE = 0.024, z = -2230, p = 0.026, Figure S9B). To explore this effect, we then performed separate logistic mixed effect regressions for each group and each stimulation type (stimulation after first-step choice, and stimulation at outcome).

In the ChR2 group, stimulating dopamine neurons after the first-step choice was reinforcing; it significantly increased the probability of repeating that choice on the next trial (β = 0.091, SE = 0.035, z = 2.645, p=0.008, mixed effect logistic regression, Figure 5F). This stimulation did not significantly interact with either the transition or outcome in its effect on next trial choice (all p>0.069). This is in line with the ICSS result (Figure 5B) and previous reports^5,9,41–44^ showing that dopamine activation promotes repeating actions.

Strikingly, stimulating dopamine neurons at the time of trial outcome – where we observed large increases and decreases in dopamine following reward or omission, respectively – had no significant influence on the subsequent choice (Figure 5I); it did not reinforce the preceding first-step choice (effect of stimulation: β = 0.041, SE = 0.030, z = 1.378, p = 0.168), nor did it act like a reward by interacting with the state transition (β = -0.011, SE = 0.030, z = -0.355, p = 0.723), nor modify the effect outcomes on choices (stimulation-transition-outcome interaction: β = -0.014, SE = 0.030, z = -0.464, p = 0.643). No effect of stimulation was found in the YFP group for either of the stimulation types (all p>0.456). To evaluate the strength of this null result in the ChR2 group, we computed a Bayes factor (B = 0.048) for whether dopamine stimulation acted like a task reward or had no effect. This indicted the manipulation result provides “strong evidence” (using the classification proposed by Jeffreys^45^) against dopamine stimulation recapitulating the behavioural consequences of rewards in this task.

By contrast, while stimulation after the first step choice had no effect on the latency to initiate the next trial (Figure 5E, t(6) = 0.347, p = 0.740, 95% CI [-30.25, 40.25], Cohen’s D = 0.034), stimulation at outcome time significantly reduced the latency to initiate the next trial (Figure 5H, t(6) = 4.228, p = 0.0055, 95% CI [20.98, 78.6], Cohen’s D = 0.369). Again, there was no effect on this latency in the YFP group (Figure 5E, stimulation after choice: t(4) = 0.816, p = 0.460, 95% CI [-62.7, 114.9], Cohen’s D = 0.302; Figure 5H, stimulation at outcome time: t(4) = 1.713, p = 0.162, 95% CI [-9.87, 41.67], Cohen’s D = 0.141).

To further corroborate that the reward effects observed in the behaviour are independent of dopamine, we repeated the previous experiment, but now using either a soma-targeted anion-conducting channelrhodopsin to inhibit VTA dopamine neurons (GtACR2, N=7) or a control fluorophore (tdTomato, n=5) (Figure S9A). Our stimulation parameters (1s continuous, 5mW) were effective at negatively reinforcing an immediately preceding action in a separate 2-alternative forced choice control task (Figure S9C). Nonetheless, inhibiting dopamine neurons during the two-step task had no effect on performance (no significant effect of stimulation by opsin group either in isolation or interacting with other trial events, all p > 0.283). To corroborate this null result, we again calculated the Bayes Factor for the GtACR2 group at outcome time stimulation (B = 0.062), which indicated “strong evidence” against dopamine inhibition modulating the effects of outcome on subsequent choices.

Therefore, while optogenetic activation or inhibition of VTA dopamine neurons was sufficient, respectively, to promote or reduce the likelihood of repeating an immediately preceding action, it completely failed to recapitulate the behavioural consequences of natural rewards at outcome, despite reward delivery and omission driving the largest dopamine signals observed in the task.

#### Neural-network model reproduces experimental data

Our behavioural analyses and dopamine recordings demonstrate subjects inferred the hidden state of the reward probabilities by observing where rewards were obtained, while our optogenetic manipulations indicate these belief updates (changes in estimates of the hidden state) were not caused by dopamine. This raises several questions: How do subjects learn there is a hidden state that controls the reward probability at both ports? Where and how are beliefs about the hidden state represented and updated? How does state inference interact with the prominent RPE signals we observe in dopamine activity?

One possibility is that recurrent networks in cortex learn to infer the hidden states by predicting observations, while reinforcement learning mechanisms in basal ganglia learn the corresponding value and appropriate action. To test this hypothesis, we implemented a simple neural network model of cortex-basal ganglia circuits (Figure 6).

**Figure 6.**
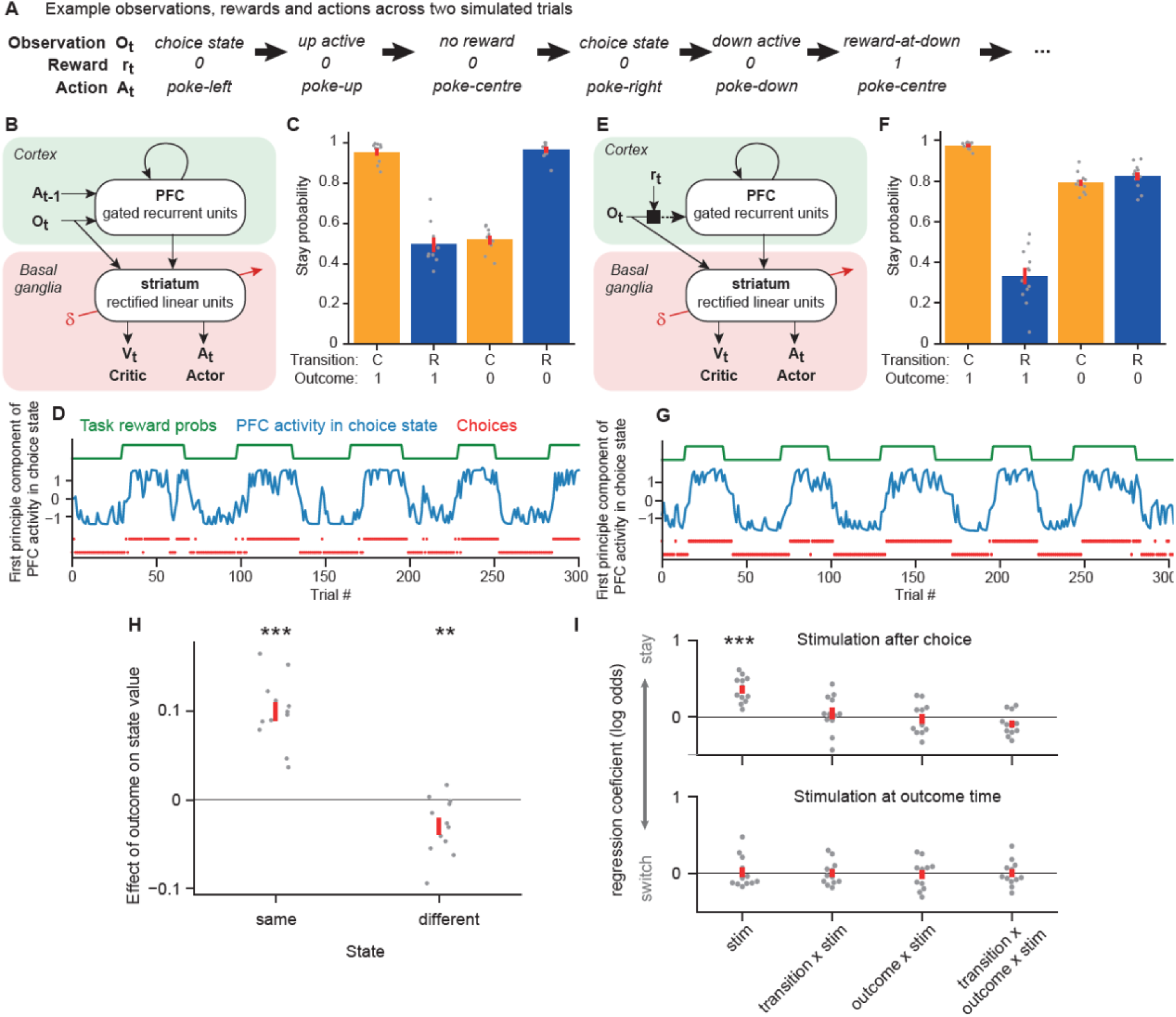
PFC-basal ganglia network model. **A)** Example observations and rewards generated by the task, and actions selected by the model, across two simulated trials comprising six time-steps. **B)** Diagram of model used in panels **C** and **D**. **C)** Stay probability analysis for behaviour generated by network model shown in **B**. **D)** Activity of the PFC network in the task’s choice state across trials (blue), projected onto the first principal component of its cross-trial variation. Both PFC activity, and the model’ choices (red), tracked the state of the task’s reward probability blocks (green). **E)** Diagram of model used in panels **F** and **G**. The only difference from the model shown in **B** was the input received by the PFC recurrent network. **F,G)** As **C**,**D** but for the model shown in **D**. **H)** Effect of trial outcome (rewarded vs non-rewarded) on the value of the second-step state where reward was received (same) and on the other second-step state (different). **I)** Effect of simulated optogenetic stimulation of dopamine neurons, either immediately after taking the first step choice (top panel) or at the time of trial outcome (bottom panel). Stimulation was modelled as modifying weights in the basal ganglia network as by a positive RPE. Figure S10 shows the analyses from panels **H** and **I** for the model shown in panel **B**.

The task was modelled as having five actions corresponding to the five nose-poke ports, and five observable states corresponding to trial events (e.g., *choice state* or *up-active*), such that completing each trial required a sequence of three actions (Figure 6A). Prefrontal cortex (PFC) was modelled as a recurrent neural network which received at each time-step an observation ***B_t_*** (the observable state) and the preceding action ***A_t-1_***. PFC activity and observations provided input to a feedforward network representing basal ganglia, comprising a single layer of rectified linear units with two output layers: a scalar estimate of the current value ***V_t_*** (i.e. expected discounted long-run reward) and a vector of action probabilities which determined the next action ***A_t_***. The PFC network was trained using gradient descent to predict the next observation given the history of observations and actions. The basal-ganglia network was trained using actor-critic reinforcement learning to estimate the value and select actions given its current input. Network weights were updated gradually across training, and held constant for the simulations analysed in Figure 6, such that changes in network activity and behaviour from trial-to-trial were mediated *only* by the changing input and recurrent activity it induced in PFC.

PFC activity tracked the hidden state of the reward probabilities across trials. Notably, this was true in the choice state (Figure 6D) even though the next observation in this state does not depend on the reward probabilities (as it is either *up-active* or *down-active*). However, to accurately predict trial outcomes the network must carry information provided by previous outcomes forward through time in its recurrent activity, causing it to be present throughout the trial. The model’s choices tracked the high reward probability option (Figure 6D), demonstrating that the basal ganglia network was able to use the reward probability information present in its PFC input to select appropriate actions.

Analysis of stay probabilities showed a strong interaction between transition and outcome; i.e., the model repeated choices following rewarded common transitions and non-rewarded rare transitions. This is exactly the pattern expected for an agent which infers the hidden state of the reward probabilities and has a fixed mapping from this to the first-step choice^33^ (Figure 1G). This pattern of stay probabilities can also be generated by model-based RL prospectively evaluating actions by predicting the states they will lead to^33,34^. However, that is not what is happening here: the PFC network only predicts the next observation *after* an action has been selected, and this prediction is used only for updating PFC network weights.

The model did not exhibit the asymmetric learning from reward and omission observed in the mice. We showed in Figure 1I-J that models which use Bayesian inference to track the reward probabilities exhibit this asymmetry if they treat rewarded outcomes as *distinct* observations based on where they occur (i.e., top/bottom port), but non-rewarded outcomes as the *same* observation irrespective of where they occur. We therefore asked if this mechanism could generate such asymmetry in the network model. We modified the input provided to the PFC network such that on each time step PFC received the observation gated by whether reward was received, such that on non-rewarded timesteps the input was a zero vector (Figure 6E). PFC activity and choices still tracked the task reward probabilities (Figure 6G), but now the stay probabilities recapitulated the asymmetry between rewarded and non-rewarded outcomes seen in the mice (Figure 6F). As with the explicitly Bayesian models (Figure 3C), the network model reproduced the non-local value updates we observed the dopamine signal (Figure 6H): i.e., reward in one second-step state increased the value of that state (t-test, t(11) = 9.28, p < 0.001) but also decreased the value of the other second-step state (t-test, t(11) = -3.23, p = 0.008).

Finally, we asked how optogenetic stimulation of dopamine neuron on individual trials would affect the model’s choice behaviour, assuming it acted by updating weights in the basal ganglia network as if it were a positive RPE (Figure 6I). Stimulation following first-step choice increased the probability of repeating the choice on the next trial (t-test, t(11) = 7.41, p < 0.001) as observed experimentally. Crucially, stimulation at trial outcome *had no effect on next trial choice* (t-test, all |t(11)| < 0.30, p > 0.77), again recapitulating the data. This is because RPE following an action updates the network weights to increase the probability of selecting the *same action* in the *same state* in future. Therefore, stimulation following a choice affects next trial choice, but stimulation following, for example, an *up-poke* in the *up-active* state has no effect on choosing left versus right in the next trial’s choice state.

In sum, the network model recapitulates our key experimental findings that both behaviour, and the value information that drives dopaminergic RPEs, are consistent with state inference, but dopamine does not mediate the effect of rewards on subsequent choices.

## Discussion

By recording and manipulating dopamine activity in a two-step decision task, we obtained a set of observations that together support an integrated framework for understanding reward-guided decision making. Rewards did not simply reinforce preceding choices, but rather promoted choosing the action that commonly led to the state where the reward was obtained, consistent with previous work with similar tasks^31,46^. Dopamine carried rich information about value, action and recent reward history, responding strongly and transiently to both rewards and states that predicted reward. The influence of state values on mesolimbic dopamine exhibited a key signature of a RPE^35–37^: a positive response when the state was entered followed by a negative response at trial outcome. Additionally, rewards obtained in one second-step state *negatively* influenced the dopamine response upon reaching the *other* second-step state on the subsequent trial, consistent with an inferred value update. Strikingly, however, neither optical activation nor inhibition of dopamine neurons at the time of trial outcome – when dopamine responses were maximal – influenced next trial choice, despite positive controls in the same subjects verifying the manipulations were effective.

We argue these data support an account where animals infer that a single hidden state controls reward probability at both ports, and track this state by observing were rewards are obtained. Critically, *these belief updates shape dopamine responses but are not caused by them*. Supporting this, we show that our data can be reproduced by a neural network model in which cortex infers hidden states by predicting observations and basal ganglia uses RL mediated by dopaminergic RPEs to learn appropriate actions.

### Reinforcement learning and inference in reward-guided choice

RL has provided an influential account of reward-guided decision making, in which the effect of rewards on subsequent choices is mediated by value updates driven by dopaminergic RPEs. However, our data are neither consistent with model-free nor model-based RL operating over the observable states of the task. Model free RL would cause rewards to reinforce preceding actions irrespective of whether a common or rare transition had occurred, inconsistent with subject’s choices. Model-based RL could explain the choice behaviour if rewards drive RPEs that update second-step state values, which determine next trial choice through prospective evaluation of first-step actions by predicting the states they will lead to. However, this predicts that dopamine manipulations at trial outcome would affect next trial choice, conflicting with the optogenetic results. Moreover, it does not explain how reward in one second-step state decreases the value of the other second-step state. By contrast, such non-local value updates *are* consistent with the animals understanding that a single hidden variable controls both reward probabilities. However, if animals solve the task by state inference, this raises important questions: how do they learn to track the hidden state, and what function do the observed dopaminergic RPEs serve^47^?

Our computational model suggests a possible answer. A recurrent network representing frontal cortex learned to track the hidden state of the reward probabilities by predicting observations. A feed-forward network representing basal ganglia learned values and appropriate actions (‘policies’) over the observable and inferred state features using RL, generating choices that closely resembled those of the subjects. Crucially, the short-timescale effect of rewards on subsequent choices was driven by changes in recurrent activity in PFC, not synaptic weight changes in either network. Consistent with this, two recent studies found that medial frontal cortex activity tracks the reward probabilities during probabilistic reversal learning, not just at decision or outcome time, but also throughout the inter-trial interval^48,49^. This is necessary if recurrent activity is responsible for carrying forward information about the recent reward history to guide choices.

Importantly, this simple model also reproduced our other key findings. For example, rewards obtained in one second-step state negatively updated the other state’s value; this occurs because rewards in either state update a common activity pattern in PFC tracking the reward probabilities, which in turn modifies value estimates for both states in basal ganglia. Moreover, simulating dopamine stimulation at outcome time had no effect on next trial choice, as there was no prospective evaluation of first-step actions in terms of the states they will lead to, so modifying values of the second-step states by evoking an RPE at outcome time has no effect. Nonetheless, just as in our experimental data, stimulating dopamine immediately after choice promoted repeating the choice on the next trial.

This two-process account of reward-guided decision making can help reconcile the burgeoning evidence for state inference regulating both behaviour^11,17–21^ and neural activity in cortex^18,50–56^ and the dopamine system^19,25–28^, with the long-standing literature supporting dopamine activity resembling and acting like an RPE signal^1–4^. It also helps explain previous findings that although stimulating/inhibiting dopamine following an action can bidirectionally modulate the probability of repeating the action in future^5–9^, inhibiting dopamine at outcome time can fail to block the effect of rewards on subsequent choices^5,6^, and pharmacological manipulation of dopamine signalling in reward guided decision tasks often has limited or no effect on learning^11,32,57,58^.

Our network model is related to a recent proposal that frontal cortex acts as a meta-reinforcement learning system^59^. In both models, synaptic plasticity acts on a slow timescale over task acquisition to sculpt recurrent network dynamics that generate adaptive behaviour on a fast timescale. Unlike this previous work, our model differentiates between cortex and basal ganglia, both with respect to network architecture (recurrent vs feedforward) and type of learning (unsupervised vs reinforcement). This builds on longstanding ideas that cortex implements a hierarchical predictive model of its sensory inputs^60,61^, while basal ganglia implement temporal difference RL^2,62^. It is also motivated by work in machine learning in which Markovian state representations (which integrate the observation history to track hidden states) are learned by predicting observations^63–66^, enabling RL algorithms to solve tasks where the current observation is insufficient to determine the correct action. There are also commonalities with connectionist models of learning phenomena where one stimulus changes the meaning of another, including occasion setting, configural and contextual learning^67,68^. These use hidden units between an input and output to allow modulatory interactions between stimuli^69,70^, just as in our model the hidden units in basal ganglia allow for non-linear combination of the current observation and PFC activity to determine value and action. Unlike this previous work, which modelled working memory for past stimuli through temporally extended input activity, our model learns what information about the past is needed to predict future observations, and stores this in PFC recurrent activity.

## RPE, value, reward rate, and movement

Recent work has questioned the relative influence of value and RPE on dopamine activity and striatal concentrations^4,5,38^. We observed the biphasic influences of the values of both first-step actions and second-step states, a key signature of RPE, in both calcium activity in VTA, NAc and DMS (Figure 3, S4), and in dopamine concentrations in NAc and DMS (Figure S5). This biphasic pattern was most prominent in NAc, with VTA and DMS signals both showing substantially larger initial positive modulation than later negative modulation. Intriguingly, when evaluating only the influence of the most recent outcome on second-step state value, rather than the extended history, the biphasic pattern was prominent in NAc but absent in VTA (Figure 3E). Differences between the VTA and striatal signals could reflect local modulation of terminals, for example by cholinergic neurons^71^, or alternatively the VTA signal may be influenced by calcium activity in dendrites that is partially decoupled from spiking due to somatic inhibition.

In parallel, we observed two other important modulators associated with dopamine function – recent reward rate and movement – both of which accounted for a separate portion of variance from the value of trial events. The average reward rate over recent trials had a strong positive influence on dopamine activity in all regions (Figure 4E, F, S4). Reward rate had sustained effects across the trial from before initiation to after outcome, unlike the influence of state and action values which were tightly time-locked to the corresponding behavioural event. This appears broadly consistent with theoretical proposals that tonic dopamine represents average reward rate, which acts as the opportunity cost of time in average-reward RL^72^, and reward-rate signalling may be responsible for the effect of dopamine manipulations on motivation and task engagement^32,58,73^. Correlation between reward rate and NAc dopamine concentrations is consistent with recent reports^5,38^, but that with VTA calcium is unexpected given previous reports of no correlation with VTA spikes^38^.

By contrast, the influence of movement was transient, lateralised and exhibited distinct dynamics in DMS and VTA/NAc. Specifically, DMS, but not NAc or VTA, dopamine was selectively influenced at the time of the initial choice, with increased activity in the hemisphere contralateral to the movement direction consistent with previous studies^6,40^. Intriguingly, at the time of the second movement from the lateralised choice port to the reward port, significant modulations were instead observed in NAc (with a similar pattern in VTA), but *not* in DMS, again with increased dopamine activity contralateral to the movement direction. This suggests that an interplay of dopamine dynamics across striatum might shape movement direction as animals proceed through a sequence to reward.

### Conclusions

Our findings emphasise that flexible behaviour involves two processes operating in parallel: inference about the current state of the world, and evaluation of those states and actions taken in them. The involvement of dopamine in updating values has rightly been a major focus of accounts of flexible decision making. However, in the structured environments common both in the lab and real world, it is only half the picture. Our data show that during reward guided decision making by experienced subjects, it is dopamine-independent state inference, not value learning, that mediates the effect of reward on future choices, revising standard accounts of dopamine’s involvement in flexible behaviour.

## Acknowledgements

This research was funded by Wellcome (202831/Z/16/Z: MEW; 214314/Z/18/Z: MEW, TA; WT096193AIA: TA; 225926/Z/22/Z: TA; 215198/Z/19/Z: MBP). For the purpose of open access, the author has applied a CC-BY public copyright license to any Author Accepted Manuscript version arising from this submission. We thank Marios Panayi for assistance with the mixed effects modelling, Tim Behrens for discussions about the work, and Peter Dayan and Armin Lak for comments on the manuscript.

## Author Contributions

TA and MEW conceived the project and, with MBP, designed the experiments. MBP and TA collected the data. MBP and TA analyzed the data. TA performed the network modelling. All authors wrote the manuscript.

## METHODS

### Subjects

All procedures were performed in line with the UK Animal (Scientific Procedure) Act 1986 and in accordance with the University of Oxford animal use guidelines, and were approved by the local ethical review panel at the Department of Experimental Psychology, University of Oxford. 12 dopamine transporter (DAT)-Cre heterozygous mice (7 females and 5 males) were used for the GCaMP photometry recordings, 6 WT C57BL/6 mice (3 females and 3 males) for the dLight recordings, and 12 DAT-Cre mice (YFP: 2 females and 3 males; ChR2: 4 females and 3 males) for the optogenetic activation experiment, and 12 DAT-Cre mice (tdTomato: 3 females and 2 males; GtACR2: 4 females and 3 males) for the optogenetic inhibition experiment. Mice were aged 8-16 weeks at the start of behavioural training. Animals were typically housed in groups of 2-4 throughout training and testing.

### Data and code availability

All data and code used in the manuscript will be made available in a public repository from acceptance of the manuscript.

#### Behavioural setup

The task was run in custom-built 12×12 cm operant boxes (design files at https://github.com/pyControl/hardware/tree/master/Behaviour_box_small) controlled using pyControl^74^. Five nose poke ports were located on the back wall of the boxes; a central initiation port flanked by two choice ports 4 cm to the left and right and two second-step ports located 1.6 cm above and below the central poke. The second-step ports each had a solenoid to deliver water rewards. A speaker located above the ports was used to deliver auditory stimuli. Video was acquired from an FLIR Flea 3 camera positioned above each setup using a Bonsai based workflow (https://github.com/ThomasAkam/Point_Grey_Bonsai_multi_camera_acquisition)^75^.

#### Behavioural task and training

The behavioural task was adapted from the human two-step task^34^. Each trial started with the central initiation port lighting up. Subjects initiated the trial by poking the illuminated port, which caused the choice ports to illuminate. On free-choice trials (75% of trials), both the left and right port lit up, allowing subjects to choose, while on forced-choice trials only one randomly selected choice port lit up, forcing animals to select that specific port. Poking a choice port was followed, after a 200ms delay, by the second-step port lighting up and 1s presentation of one of two auditory cues (‘second-step cue’, either a 5- or 12-kHz tone depending on whether the top or bottom second-step became active, counterbalanced across animals). Each choice port was commonly (80% trials) associated with transitioning to one second-step state (up or down) and rarely (20% trials) to the other. The transition structure was fixed for each animal across all sessions, and was counterbalanced across animals – i.e. the task had 2 possible transition structures: transition type A, where left choice commonly led to the up second-step port, and right choice commonly led to the down second-step port; and the opposite for transition type B. The second-step port only became responsive to pokes after cue offset. Poking the second-step port triggered a 200ms delay, after which a 500ms auditory cue signalled whether the trial was rewarded or not (same 5-or 12-kHz tone as the second-step cue, counterbalanced across animals, pulsed with at 10Hz on rewarded trials, white noise on unrewarded trials). Reward was delivered at the offset of this cue. To ensure mice knew when they had made a nose poke, a click sound was presented whenever the subject poked a port that was active (e.g., the initiation port during the initiation state).

Reward probabilities for the two second-step ports changed in blocks. In non-neutral blocks, one second-step port had 80% reward probability and the other 20%, while in neutral blocks both second-step ports were rewarded with 50% probability. Block transitions from non-neutral blocks were triggered 5 to 15 trials after mice crossed a threshold of 75% “correct” choices (i.e., choosing the higher reward probability), computed as the exponential moving average with time constant of 8 free choice trials. Transitions from neutral blocks were triggered after a 20-30 trials. An inter-trial interval of 2-4 s duration started once the subject remained out of the second-step port for 250ms after the trial outcome.

##### Training

Behavioural training took 4-6 weeks. Animals were put on water restriction 48 hours before starting training, and received 1 hour ad lib water access in their home cage 24 hours before starting training. On training days (typically 6 days/week) animals usually received all their water from the task, but were topped up outside the task as necessary to maintain a body weight >85% of their pre restriction baseline. On days off from training mice received 1 hour ad lib water access in their home cage. Water reward size was decreased from 15ul to 4ul across training in order to increase the number of trials performed and blocks experienced on each session.

Training consisted of multiple stages with increasing complexity (Table 1). At the first training stage 1.1, only the second-step ports were exposed, with all other ports covered. Second-step ports were illuminated in a pseudorandom order with inter-trial interval of 2-4 seconds. Poking an illuminated port delivered reward with 100% probability, with no auditory cues. When animals obtained >50 rewards on this stage, transitioned to stage 1.2, where the auditory cues for second-step state and reward were introduced. Once animals obtaining >70 rewards in a session, they were switched to stage 2 on the next session. At stage 2, the choice ports were introduced, but all trials were forced choice, such that only one choice port lit up on each trial. Mice were switched to stage 3 when they obtained >70 rewards on a single session. At stage 3, the initiation poke was introduced, and when they obtained >70 rewards on a single session, they transitioned to stage 4 on the next training session. Stage 4 consisted of multiple substages where progressively more free choice trials were introduced, and the reward probabilities gradually changed until reaching the final task parameters. Mice were transitioned to next substage after two training sessions of 45min or a single 90min session, until they reached substage 4.6. Subjects were only transition to the final stage (full task) when they completed at least 5 blocks in a single session.

**Table 1.** Description of training stages and task contingencies.

| Stage | Start poke | Choice pokes | Reward pokes | Task Structure |
| --- | --- | --- | --- | --- |
| 1.1 |  |  | X | Reward probability 100% |
| 1.2 |  |  |  | Introduce auditory cues<br>Reward probability 100% |
| 2 |  | X | X | Transition probability 80/20%<br>Reward probability 100% |
| 3 | X | X | X | Transition probability 80/20%<br>Reward probability 100% |
|  | x | x | x | Start introducing free choice trials<br>and reward probabilities |
| Substage | Free choice trials | Non-neutral blocks<br>reward probabilities<br>(high / low) | Neutral block<br>reward probability | Block change<br>(independent / dependent on<br>behaviour) |
| 4.1 | 0% | 0.9/0.7 | 0.8 | Independent – 20-30 trials blocks |
| 4.2 | 25% | 0.9/0.5 | 0.7 | Independent – 20-30 trials blocks |
| 4.3 | 25% | 0.9/0.3 | 0.6 | Independent – 20-30 trials blocks |
| 4.4 | 25% | 0.9/0.1 | 0.5 | Independent – 20-30 trials blocks |
| 4.5 | 50% | 0.9/0.1 | 0.5 | Dependent |
| 4.6 | 75% | 0.9/0.1 | 0.5 | Dependent |
| 4.7 – Final | 75% | 0.8/0.2 | 0.5 | Dependent |

#### Surgery

Mice were anaesthetised with isoflurane (3% induction, 0.5-1% maintenance), and injected with buprenorphine (0.08 mg/kg), meloxicam (5 mg/kg) and glucose saline (0.5ml). Marcaine (max 2 mg/kg) was injected into the scalp and before placing mice into the stereotaxic frame. Mice were maintained at ∼37 degrees with rectal probe and heating blanket (Harvard Apparatus). Surgery proceeded as described below for the different experiments. After surgery, mice were given additional doses of meloxicam each day for 3 days after surgery, and were monitored carefully for 7 days post-surgery.

##### GCaMP photometry

Mice were intracranially injected with 1ul of saline containing 1:10 dilution of AAV1.Syn.Flex.GCaMP6f.WPRE.SV40 (titre: 6.22 × 10^12^ vg/ml, Penn Vector Core) and 1:20 dilution of AAV1.CAG.Flex.tdTomato.WPRE.bGH (AllenInstitute864) (titre: 1.535 × 10^12^ vg/ml, Penn Vector Core) at 2nl/s in VTA (antero-posterior (AP): -3.3, medio-lateral (ML): ±0.4, dorso-ventral (DV): -4.3 from bregma) in one hemisphere for mesolimbic dopamine recordings in VTA and NAc and VTA/SNc (AP: -3.1, ML: ±0.9, DV: -4.2 from bregma) in the other hemisphere for DMS recordings. Three 200 µm diameter ceramic optic fibres were implanted chronically in each animal in VTA (AP: -3.3, ML: ±0.4, DV: -4.3 from bregma) and NAc (AP: +1.4, ML: ±0.8, DV: -4.1 from bregma) in the same hemisphere, and DMS (AP: +0.5, ML: ±1.5-1.7, DV: -2.6 from bregma) in the contralateral hemisphere.

##### dLight photometry

Mice were intracranially injected with 500 nl of saline containing 1:5 dilution of pAAV5-CAG-dLight1.1 (titre: 1.4 × 10^12^ vg/ml, addgene) and 1:5 dilution of pssAAV-2/5-hSyn1-chI-tdTomato-WPRE-SV40p(A) (titre: 4.9 × 10^11^ vg/ml, ETH Zurich) at 2nl/s in NAc (AP: +1.4, ML: ±0.8, DV: -4.1 from bregma) and DMS (AP: +0.5, ML: ±1.5/1.7, DV: -2.6 from bregma) in opposite hemispheres. Two 200 µm diameter ceramic optic fibres were implanted chronically in the injection sites.

##### Optogenetic manipulation

For optical activation experiments, mice were injected bilaterally either with 500 nl/hemisphere of saline containing either AAV2-EF1a-DIO-EYFP (titre: >1 × 10^12^ vg/ml, UNC Vector Core) (YFP group) or rAAV2/Ef1a-DIO-hchR2(E123t/T159C)-EYFP (titre: 5.2 × 10^12^ vg/ml, UNC Vector Core) (ChR2 group) at 2 nl/s in VTA (AP: -3.3, ML: ±0.4, DV: -4.3 from bregma). For optical inhibition experiments, mice were injected bilaterally with 500 nl/hemisphere of saline containing 1:10 dilution of either ssAAV-1/2-CAG-dlox-tdTomato(rev)-dlox-WPRE-bGHp(A) (titre: 7.9 × 1012 vg/ml, ETH Zurich) (tdTomato group) or AAV1-hSyn1-SIO-stGtACR2-FusionRed (titre: 1.9 × 1013 vg/ml, addgene) (GtACR2 group) at 2 nl/s in VTA (AP: -3.3, ML: ±0.4, DV: -4.3 from bregma). For both sets of experiments, two 200 µm diameter ceramic optic fibres were implanted chronically targeting the injection sites at a 10-degree angle.

#### Histology

Mice were terminally anesthetised with sodium pentobarbital and transcranial perfused with saline then 4% PFA solution. 50 µm coronal brain slices were cut, covering striatum and VTA, and immunostained with anti-GFP and anti-TH primary antibodies, and 488 and Cy5 secondary antibodies. For the animals used on the optogenetic inhibition experiment, only anti-TH and Cy5 primary and secondary antibodies, respectively, were used. An Olympus FV3000 microscope was used to image the slices.

#### Photometry recordings

Dopamine calcium activity (GCaMP) and release (dLight) were recorded at 130Hz sampling rate using pyPhotometry^76^. The optical system comprised a 465 nm and a 560 nm LED, a 5 port minicube and fibre optic rotary joint (Doric Lenses), and two Newport 2151 photoreceivers. Time division illumination with background subtraction was used to prevent crosstalk between fluorophores due to the overlap of their emission spectra, and changes in ambient light from affecting the signal. Synchronisation pulses from pyControl onto a digital input of the pyPhotometry board were used to synchronise the photometry signal with behaviour^76^.

Photometry signals were pre-processed using custom Python code. A median filter (width 5 samples) was first used to remove any spikes due to electrical noise picked up by the photodetectors. Afterwards, a 5Hz low-pass filter was used to denoise the signal. To motion-correct the signals, we band-passed the denoised signals between 0.001 Hz and 5 Hz, and used linear regression to predict the GCaMP or dLight signal using the control fluorophore (tdTomato) signal. The predicted signal due to motion was subtracted from the denoised signal. To correct for bleaching of the fibre and fluorophores, signals were detrended using a double exponential fit to capture the temporal dynamics of bleaching: a first fast decay and a second slower one (Figure S11). Finally, the pre-processed dopamine signal was z-scored for each session to allow comparison across sessions and animals with different signal intensities.

For the GCaMP data, we recorded from 12, 11 and 12 mice in VTA, NAc and DMS, respectively (in one animal, the fibre targeting NAc did not exhibit any GCaMP modulation, which later histological analysis confirmed was caused by the fibre being in the anterior commissure). In DMS, 2 animals – the two with most medial coordinates – were excluded from the main analyses of the effects of reward on subsequent dopamine activity as they presented a negative modulation to reward (Figure S3).

For the dLight data, we recorded from 5 and 6 mice in NAc and DMS, respectively (1 subject in NAc did not present any dLight modulation, posterior histological analysis confirmed that the fibre was misplaced into the ventricle).

Sessions in which there were large artifacts (large step change in recorded signals) introduced through a malfunctioning of the rotary joint or disconnection of the patch cord from the fibre, or where there was a complete loss of signal on one of the channels due to discharged battery during recording, were excluded. A total of 46 sessions (∼9% of the total) were removed from the analyses.

#### Optogenetic activation

VTA dopamine neurons were stimulated bilaterally using two 465 nm LEDs (Plexon Plexbright) connected to 200 µm 0.66 NA optic fiber patch cords. All stimulation experiments used stimulation parameters of 5 pulses at 25 Hz, 25% duty cycle, ∼8-10mW optical power at the fibre tip.

First, as a positive control, we performed an intracranial self-stimulation (ICSS) assay in which mice were presented with either 4 or 2 nose poke ports, one of which triggered optical stimulation when poked. A minimum 1-second delay was imposed between stimulations. Mice were tested on ICSS on 40-60min sessions for 4 days.

We then tested the effect of optogenetic activation during the two-step task. On each stimulation session, we either stimulated (i) 200ms after the first-step choice at the time of second-step cue onset, or (ii) at outcome-cue onset. Stimulation occurred on 25% of the trials, under the constraints that (i) the trial after stimulation was always a free choice trial, and (ii) there were always at least 2 non-stimulated trials after each stimulation. The stimulation sessions were interspersed with baseline no stimulation sessions (data not shown). The timing of stimulation was fixed within a session, with the session order counterbalanced across animals (e.g., no stimulation session → second-step cue stimulation session → outcome stimulation session → no stimulation session → outcome stimulation session → second-step cue stimulation session, etc).

#### Optogenetic inhibition

VTA dopamine neurons were inhibited bilaterally using two 465 nm LEDs (Plexon Plexbright) connected to 200 µm 0.66 NA optic fiber patch cords. All inhibition experiments used stimulation parameters of 1s continuous light at ∼5mW optical power at the fibre tip.

We first tested the effect of optogenetic inhibition on the two-step task. As in the stimulation experiment, on each inhibition session, we either inhibited (i) 200ms after the first-step choice at the time of second-step cue onset, or (ii) at outcome-cue onset; baseline sessions with no stimulation interspersed with the stimulation sessions (data not shown). Inhibition occurred on 25% of the trials under the same constraints as in the optogenetic activation experiment (see above).

As a positive control, we then performed a 2-alternative forced choice bias assay. Mice were presented with 3 ports: a central initiation port, and left and right choice ports. Mice initiated each trial by poking the illuminated centre port. This either triggered both the left and right ports to light up (free choice trials, 50% of total) or just one choice port to illuminate. Poking an illuminated choice port led, after a 200 ms delay, to 500 ms presentation of an outcome cue (5-or 12-kHz tone – left or right frequency tone counterbalanced across animals – pulsed with at 10Hz on rewarded trials, white noise on unrewarded trials), after which, on rewarded trials, reward was delivered. The reward probability associated with choice of either the left and right ports was fixed at 50% throughout. After 3 days of training without optical stimulation and once animals showed a consistent bias towards one of the choice ports for at least 2 consecutive days (termed the animal’s preferred choice), stimulation sessions commenced. In these, 1s continuous light stimulation was delivered coincident with the outcome cue on any trial when the preferred choice port was selected (on both free-and forced-choice trials). After 4 days, the light stimulation was then switched to be paired with selection of the other choice port for 4 more days. Each day animals were tested on a single 60min session.

#### Analysis

All behavioural and photometry analysis was performed using custom Python and R code.

##### Behavioural logistic regression model

The logistic regression analysis shown in Figure 1E predicted repeating choices (or “staying”) as a function of the subsequent trial events, considering only free choice trials, implemented as a mixed effects model using the afex package^77^ in R programming language. The model formula was:

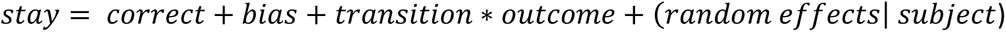

For the analysis of two-step optogenetic manipulation (Figure 5F, I, S9), we added stimulation (stim) and group and its interaction with trials events as additional predictors, giving the formula:

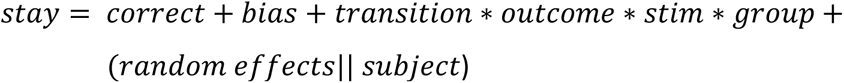

We used orthogonal sum-to-zero contrasts, and likelihood ratio test to calculate p-values. The maximal random effect structure^78,79^ with subject as a grouping factor was used.

The predictors were coded as:

- *Correct*: 3 level categorical variable indicating whether the previous choice was correct, incorrect or in a neutral block.
- *Bias*: Binary categorical variable explaining whether previous choice was left or right.
- *Outcome*: Binary categorical variable indicating whether the previous trial was rewarded or not.
- *Transition*: Binary categorical variable indicating whether previous trial had a common or a rare transition.
- *Stimulation*: 3 level categorical variable indicating whether previous trial was non-stimulated, stimulated after choice, or stimulated at outcome time.
- *Group:* Binary categorical variable indicating whether the data is from control or manipulation subject. YFP and ChR2 for the stimulation experiment, and tdtomato and GtACR2 for the inhibition experiment.

For the optogenetic manipulation analysis, we performed a single regression for the activation experiment including both experimental groups (YFP & ChR2) and both stimulation times (after choice & outcome). We did the same for the inhibition experiment (tdtomato & GtACR2). To ensure stimulated and non-stimulated trials had matching histories, we only included trials where stimulation could have potentially been delivered, i.e., excluding the 2 trials following each stimulation where stimulation was never delivered. As a follow-up analysis, we performed separate regressions per group (YFP, ChR2, tdtomato & GtACR2) and stimulation time (after choice & outcome).

To test the strength of evidence of our null results, we performed Bayes Factor calculation using R as:

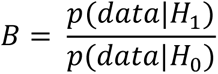

We defined the data likelihood as a normal distribution with the mean and standard deviation of the transition x stimulation regression coefficient. For the optogenetic activation experiment, the alternative hypothesis, *H*_1_, was that dopamine activation acted like a natural reward, defined as a uniform distribution between 0 and the transition x outcome regression coefficient. For the optogenetic inhibition experiment, *H*_1_ was that dopamine inhibition *reduced* the effects of natural rewards, defined as a uniform distribution between 0 and minus the transition x outcome regression coefficient. Finally, the null hypothesis, *H*_0_, was set to 0. We used the classification proposed by Jeffreys^45^ to assess the strength of evidence for the alternative or null hypothesis.

The lagged logistic regression analysis (Figure 1F, H) assessed how subjects’ choices were affected by the history of trial events over the last 12 trials. The regression predicted subjects’ probability of choosing left, using the following predictors at lags 1, 2, 3-4, 5-8, 9-12 (where lag 3-4 etc. means the sum of the individual trial predictors over the specified range of lags).

- *Common Transition*: Rewarded at lag *n*: +0.5/-0.5 if the *n*’th previous trial was a left/right choice followed by a common transition and reward, 0 otherwise.
- *Rare Transition*: Rewarded at lag *n*: +0.5/-0.5 if the *n*’th previous trial was a left/right choice followed by a rare transition and reward, 0 otherwise.
- *Common Transition*: Unrewarded at lag *n*: +0.5/-0.5 if the *n*’th previous trial was a left/right choice followed by a common transition and no reward, 0 otherwise.
- *Rare Transition*: Unrewarded at lag *n*: +0.5/-0.5 if the *n*’th previous trial was a left/right choice followed by a rare transition and no reward, 0 otherwise.

The lagged regression was fitted separately for each subject as a fixed effects model. Cross subjects mean and standard error for each predictor coefficient were plotted, significance of coefficients was assessed using a two-sided t-test comparing the distribution of the individual subjects’ coefficients against zero, and performing Bonferroni multiple comparison correction.

##### Single-strategy models

We evaluated goodness-of-fit to subjects’ choices for a set of different reinforcement learning (RL) agents created by combining one or more of the following learning strategies:

###### Model-free

The model-free strategy updated value *Q*_*MF*_(*C*) of the chosen action and value *V*(*s*) of second-step state as:

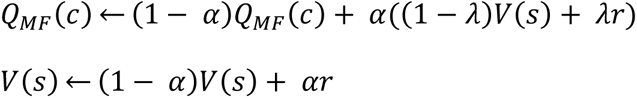

Where *α* is the learning rate, *λ* is the eligibility trace parameter and *r* is the outcome.

A variation of this model was the asymmetric model-free strategy which included different learning rates for positive and negative outcomes, and included forgetting of the non-experienced states, so both the non-experienced action and second-step state decayed towards neutral value (0.5).

###### Model-based

The model-based strategy updated value *V*(*s*) of the second-step reached, and both first-step action values *Q*_*MB*_(*a*) as:

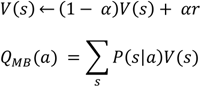

Where *α* is the learning rate, *λ* is the outcome, *λ*(*s*|*a*) is the transition probability of reaching second-step *s* after taking action *a* (i.e. 0.8 or 0.2).

A variation of this model was the asymmetric model-based strategy which included different learning rates for positive and negative outcomes, and included forgetting of the non-experienced second-step state which decayed towards neutral value (0.5). In addition, we tested another model in which the forgetting rate decayed to 0 (Asymmetric model-based + forget to 0).

###### Bayesian inference

The Bayesian inference strategy treated the task as having a binary hidden state ℎ ∈ {*up_good_down_good*} which determined the reward probabilities for the two second-step states as:

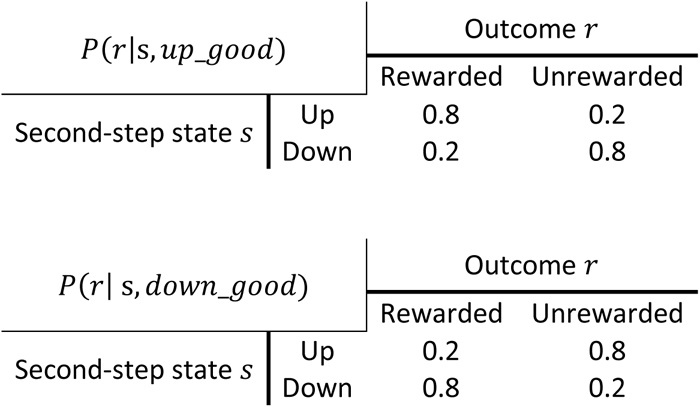

The strategy maintained an estimate *P*(*up*_*good*) tracking the probability the task was in the *up*_*good* hidden state, updated following each trial’s outcome using Bayes rule as:

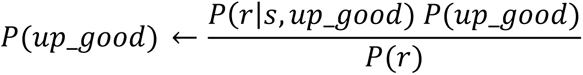

Where *P*(*r*) = *P*(*r*|*s, up*_*good*) *P*(*up*_*good*) + *P*(*r*|*s, down*_*good*) *P*(*down*_*good*)

*P*(*up*_*good*)was also updated each trial based on the probability a reversal occurred as:

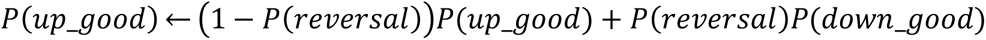

Where *P*(*reversal*) is the probability a block reversal occurred.

The value of second-step states and first-step actions was determined by *P*(*up*_*good*) as:

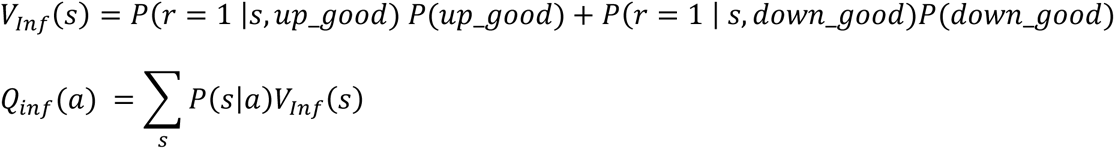

Note that though the equation relating first-step action values *Q*(*a*) to second-step state values *V*(*s*) is the same for the inference and model-based RL strategies, the mechanistic interpretation is different: For the inference strategy the action values are assumed to have been learned gradually over task acquisition using temporal difference RL operating over a state representation combining the observable state *s* and belief state *P*(*up*_*good*). This learning process was modelled explicitly in the network model (Figure 6), which generated choice behaviour similar to the inference model (Figure 1). For the model-based RL strategy the first-step action values are assumed to be computed online by predicting the states the actions will lead to.

We also used a variant of the inference strategy designed to capture the asymmetric influence of reward and omission on subject’s choices. This treated rewards in each second-step state as different observations but treated reward omission at the up and down second-step states as the same observation. The corresponding Bayesian update to *P*(*up*_*good*) given each trial’s outcome was given by:

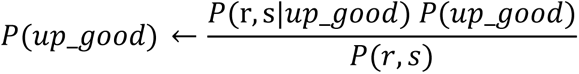

Where *P*(*r*,*s*)= *P*(*r*,*s*|*up*_*good*) *P*(*up*_*good*)+ *P*(*r*,*s down*_*good*) *P*(*down*_*good*)

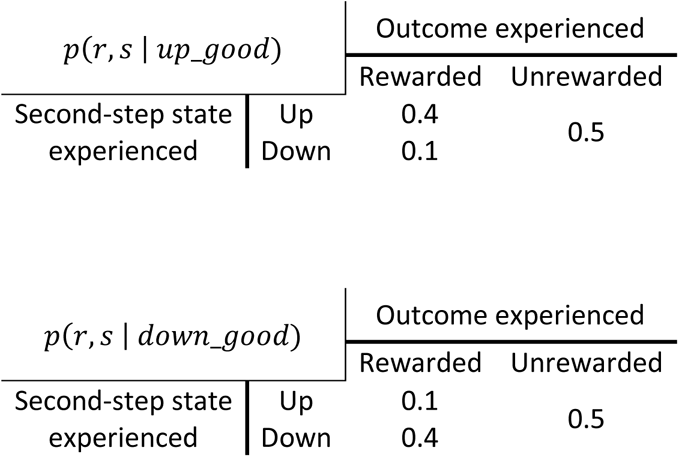

State and action values (*V*_*inf*_(*s*) and *Q*_*inf*_(*a*)) for the asymmetric inference strategy were computed as for the standard inference strategy.

###### Combined action values

A set of different candidate models were created by combining one of more of the above strategies in a weighted sum with (optionally) bias and perseveration parameters, to give net action values:

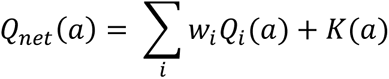

Where *W_i_* is the weight assigned to strategy *i* whose first step action values are given by *Q_i_*(*a*), and *K*(*a*) is the modifier to the value of first-step action *a* due to any bias or perseveration terms included in the model. In models where bias was included, this increased the value of the left action by an amount determined by a bias strength parameter on all trials. In models where perseveration was included, this increased the value of the first step action chosen on the previous trial by an amount determined by a perseveration strength parameter.

The combined action values determined choice probabilities via a softmax decision rule as:

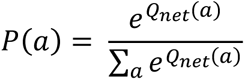

##### Model fitting and comparison

We generated a total of 53 different individual models from the following classes:

###### Model-free (MF)

These models used only the model-free strategy, and varied with respect to whether they had asymmetric learning rates for positive and negative outcomes (together with forgetting towards neutral), and whether they included perseveration or multi-trial perseveration and/or bias.

###### Model-based (MB)

These models used only the model-based strategy, and varied with respect to whether they had asymmetric learning rates for positive and negative outcomes (together with forgetting to either neutral or 0), and whether they included perseveration or multi-trial perseveration and/or bias.

###### Hybrid (MF + MB)

These models used both the model-based and model-free strategy, and varied with respect to whether they had asymmetric learning rates for positive and negative outcomes (together with forgetting towards neutral), and whether they included perseveration or multi-trial perseveration and/or bias.

###### Bayesian inference

These models used the Bayesian inference strategy, and varied with respect to whether they included an asymmetric updating based on the outcome received, and whether they included perseveration or multi-trial perseveration and/or bias.

Bias increased the value of the left action by an amount determined by a bias strength parameter. Perseveration increased the value of the first-step action chosen on the previous trial by an amount determined by a perseveration strength parameter; in the case of multi-trial perseveration, an exponential moving average of previous choices was used rather than just the previous choice, with a time constant determined by the alpha multi-trial perseveration parameter.

Each model was fit separately to data from each subject using maximum likelihood. The optimisation was repeated 30 times starting with randomised initial parameter values drawn from a Beta distribution (*α*=2, β=2) for unit range parameters, Gamma distribution (*α*=2, β=0.4) for positive range parameters, and Normal distribution (σ=5) for unconstrained parameters, and the best of these fits was used. Model comparison was done using both Bayesian Information criterion (Figure S1A, Table 1, 2), and cross-validated log likelihood using 10 folds (Figure S1B).

To compare data simulated from the single-strategy models with real data (Figure S1G-J), we simulated the same amount of sessions for each animal (28.9 ± 1.6 sessions, mean ± standard deviation) of their average number of trials per session (351.6 ± 93.0 trials, mean ± standard deviation), using parameters values from each animal’s fits.

#### Mixture-of-strategies model

We created a mixture-of-strategies models which contained the different behavioural strategies from the single-strategy models (model-free RL, model-based RL and Bayesian inference) as components (Figure S1C-F, Table 3). All 3 components included asymmetric updating from rewards and omissions.

This mixture-of-strategies model combined the action values of the 3 strategies in a weighted sum with bias and multi-trial perseveration to give net action values as:

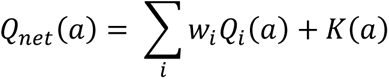

Where *W_i_* is the weight assigned to strategy *i* whose first step action values are given by *Q_i_*(*a*), and *K*(*a*) is the modifier to the value of first-step action *a* due to bias and multi-trial perseveration. The combined action values determined choice probabilities via a softmax decision rule like in the single-strategy models.

This model was fit separately to data from each subject using maximum a posteriori (MAP) probability, with priors: Beta distribution (*α*=2, β=2) for unit range parameters, Gamma distribution (*α*=2, β=0.4) for positive range parameters, and Normal distribution (σ=5) for unconstrained parameters. The optimisation was repeated 50 times starting with randomised initial parameter values drawn from the prior distributions.

To test whether behaviour generated by the model-based and inference strategies could be differentiated, we fit the mixture-of-strategies model to data simulated from each single-strategy model using parameters fit to subjects’ data (Figure S1C,D).

#### Photometry analysis

Photometry signals were aligned across trials by linearly time-warping the signal at the two points in the trial where timings were determined by subject behaviour; between initiation and choice, and between the second-step port illuminating and being poked (Figure 2B). Activity at other time periods was not warped.

For the analyses presented in Figures 4, 5, S4-8, we used Lasso linear regression to predict trial-by-trial dopamine activity at each timepoint in a trial as:

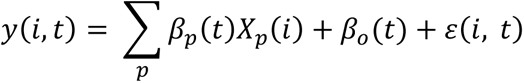

where *y*(*i, t*) is the calcium z-scored activity on trial *i* at timepoint *t*, β_*p*_(*t*) is the weight for predictor *p* at timepoint *t, X*_*p*_(*i*) is the value of the predictor *p* on trial *i*, β_*o*_(*t*) is the intercept at timepoint *t*, and *ε*(*i, t*) is the residual unexplained variance.

The linear regression was fit separately for each subject to obtain the coefficient time-courses β_*p*_(*t*). The penalty used for the Lasso regularisation was found for each individual regression through cross-validation. When regularisation was used, predictors were standardised by centring the mean at 0 and scaling to variance of 1. For each predictor we plot the mean and standard error across subjects. Statistical significance of each predictor at each timepoint was assessed using a t test comparing the distribution of coefficients across subjects with zero, with Benjamini-Hochberg correction for comparison of multiple timepoints. Effect sizes were computed using Cohen’s d at each timepoint as:

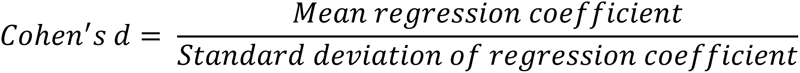

The linear regressions in Figures 3 & S4-5 used the following predictors:

- *Reward*: +0.5 if current trial is rewarded, -0.5 otherwise.
- *Previous reward*: +0.5 if previous trial is rewarded, -0.5 otherwise.
- *Good Second-Step*: +0.5 if the second step reached on the current trial has high reward probability, -0.5 if low reward probability, 0 if neutral block.
- *Previous Good Second-Step*: +0.5 if the second step reached on the previous trial has high reward probability, -0.5 if low reward probability, 0 if neutral block.
- *Correct Choice*: +0.5 if subject current trial choice commonly leads to the high reward probability second-step port, -0.5 if to the low reward probability second-step port, 0 if neutral block.
- *Repeat Choice*: +0.5 if same choice as previous choice, -0.5 if different choice to previous trial.
- *Direct Reinforcement Action Value Update*: +0.5/-0.5 if current choice is the same as the previous choice and the previous trial was rewarded/not-rewarded, 0 if different choice from previous trial.
- *Inferred Action Value Update*: +0.5/-0.5 if current choice commonly leads to the previous second-step when it was rewarded/not-rewarded, -0.5/+0.5 if current choice rarely leads to the previous second-step when it was rewarded/not-rewarded.
- *Previous Reward, Same Second-Step*: +0.5 if current second-step is the same as in the previous trial and previous trial was rewarded, -0.5 if same second-step and previous trial was unrewarded, 0 if current second-step different from second-step on the previous trial.
- *Previous Reward, Different Second-Step*: +0.5 if current second-step is different from previous second-step and previous trial was rewarded, -0.5 if different second-step and previous second-step was unrewarded, 0 if current second-step is the same as the second-step on the previous trial.
- *Common Transition*: +0.5 if a common transition occurs on the current trial, -0.5 if a rare transition.
- *Forced choice*: +0.5 if a forced choice trial occurs on the current trial, -0.5 if free choice.
- *Reward Rate*: exponential moving average of the recent reward rate (tau=10 trials).
- *Contralateral Choice*: +0.5 if current choice is in the contralateral side from the recording site, -0.5 if current choice is in the ipsilateral side.
- *Up Second-Step:* +0.5 if current second-step is up, -0.5 if current second-step is down.

The linear regression in Figure S6C, D use the same predictors as above (Figures 3 & S4-5) except predictors *Same Second-Step, Previous Reward, Different Second-Step* and *Inferred Action Value Update*, which were replaced with predictors split by outcome as:

- *Same vs Different Second-Step, Previously Rewarded*: +0.5/-0.5 if current second-step is the same/different as in the previous trial and the previous trial was rewarded, 0 if the previous trial was not rewarded.
- *Same vs Different Second-Step, Previously Non-Rewarded*: +0.5/-0.5 if current second-step is the same/different as in the previous trial and the previous trial was not rewarded, 0 if the previous trial was rewarded.
- *Inferred Action Value Update from Rewarded Trials*: +0.5/-0.5 if current choice commonly/rarely leads to the previous second-step and the previous trial was rewarded, 0 if the previous trial was not rewarded.
- *Inferred Action Value Update from Unrewarded Trials*: -0.5/+0.5 if current choice commonly/rarely leads to the previous second-step and the previous trial was not rewarded, 0 if the previous trial was rewarded.

The linear regressions used in Figures 4 & S7-8 used the above-described *Reward*, Previous Reward, *Reward Rate, Contralateral choice, Up second-step, Common transition, and forced choice* regressors with the following additional regressors:

- *Second-Step Value:* The value *V*_*inf*_(*s*) of the second-step reached on the current trial from the Asymmetric Bayesian Inference model.
- *Chosen Action Value*: The value *Q*_*inf*_(*c*) of the first-step action chosen on the current trial from the Asymmetric Bayesian Inference model.

Finally, the lagged photometry regression in Figure S6 predicted dopamine response (500ms at the end of the second-step cue, baseline subtracted using the 500ms before choice) to the second-step cue as a function of the extended history of trials over the previous 12 trials. No regularization was used in this linear regression. The regressors included the above-described: *Previous Reward, Same Second-Step; Previous Reward, Different Second-Step; Direct Reinforcement Action Value Update; and Inferred Action Value Update* regressors at different lag *n, with the following additional regressors to correct for correlations in the signal:*

- *Same vs Different Second-Step*: +0.5/-0.5 if current second-step is the same/different as in the *n*’th previous trial.
- *Reward on trial* -1 (not lagged): +0.5 if previous trial is rewarded, -0.5 otherwise.

## Neural Network model

For the neural network modelling (Figure 6), the task was represented as having 5 observable states; *choice state, up-active, down-active, reward-at-up, reward-at-down, no-reward,* and 5 actions corresponding to the 5 ports: *poke-left, poke-right, poke-up, poke-down and poke- centre.* Completing each trial therefore required a sequence of at least 3 states and actions (e.g., *choice-state, poke-left →up-active, poke-up →reward-at-up, poke-centre),* but could take more steps if the agent choose actions that were inactive in the current state (e.g., *poke-left* in the *up-active* state).

The neural network model consisted of a recurrent network representing prefrontal cortex (PFC) and a feedforward network representing basal ganglia (BG), implemented using the Keras Tensorflow API (https://keras.io). The PFC network was a single fully connected layer of 16 Gated Recurrent Units (GRUs)^80^. In the version of the model shown in figure 6 B-D, the

PFC network received as input on each time-step an observation ***O***_*t*_ (the observable task state) and the preceding action ***A_t-1_***, both coded as 1-hot vectors. In the version of the model shown in figure 6 E-I, the PFC network received as input a vector 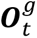 or negative which was the observation ***O***_*t*_ gated by whether reward was received on that time-step: On rewarded time-steps the input was a 1-hot vector indicating the observation 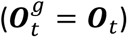, while on non-rewarded time-steps the input was a 0 vector 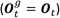.

The BG network received as input the observation ***O***_*t*_ and the activity of the PFC network units. It comprised a layer of 10 rectified linear units with two outputs: a scalar-valued linear output for the estimated value *V*_*t*_ (i.e., the expected discounted future reward from the current time-step), and a vector-valued softmax output for the policy (i.e. the probability of choosing each of the 5 actions on the next time-step).

The model was trained using episodes which terminated after 100 trials or 600 time-steps (whichever occurred first), with network weights updated between episodes. For the version of the model used in Figure 6B-D the PFC network was trained to predict the observation ***O***_*t*_ given the preceding observations and actions. For the version of the model used in Figure 6E-I, the PFC network was trained to predict the reward-gated observation 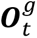 which it received as input, given this input on preceding time-steps. In both cases, PFC network weights were updated using gradient descent with backpropagation through time, with a mean-squared-error cost function, using the Adam optimiser^81^ with learning rate = 0.01. The BG network was trained using the Advantage Actor-Critic (A2C) reinforcement learning algorithm^82^. Hyperparamters for training the BG network were: learning rate = 0.05, discount factor = 0.9, entropy loss weight = 0.05.

For each version of the model, we performed 12 simulation runs, each of 500 episodes, using different random seeds. We used these runs as the experimental unit for statistical analysis (i.e., the equivalent of subjects in the animal experiments). Data from the last 10 episodes of each run were used for analyses. We excluded any runs that did not obtain reward above chance level in the last 10 episodes, excluding 2 runs of the model version shown in Figure 6B-D and no runs of the model variant shown in Figure 6E-I.

To visualise how the activity of PFC units tracked the reward probability blocks, we took the activity of the PFC units in the task’s *choice state* on each trial of an episode, yielding an activity matrix of shape [*n_units, n_trials*]. We used PCA to find the first principal component of the activities’ variation across trials (a vector of weights over units), then projected the activity matrix onto this, giving the time series across trials shown in (Figure 6D,G).

To evaluate how rewards modified the value of the second-step states (Figure 6I), we used the model to evaluate both the second-step state that was actually reached on each trial, and the value the other second-step state would have if it had been reached. We then computed how the trial outcome (reward vs omission) changed the value of the second-step state where it was received, and the other second-step state.

To simulate the effects of optogenetic stimulation of dopamine neurons (Figure 6I), we randomly selected 25% of trials and for each of these trials computed the update to the BG network weights that would be induced by a positive reward prediction error occurring either following the choice action (Choice time stimulation), or following the second-step action (Outcome time stimulation). We evaluated how these weight updates affected behaviour using linear regression to model the probability of repeating the choice on the next trial (stay probability) as a function of the transition, outcome and whether the trial was stimulated or not.

Complete code used to simulate the model and generate figure 6 is available at: https://github.com/ThomasAkam/PFC-BG_model.

## SUPPLEMENTARY FIGURES AND TABLES

**Supplementary Figure 1.**
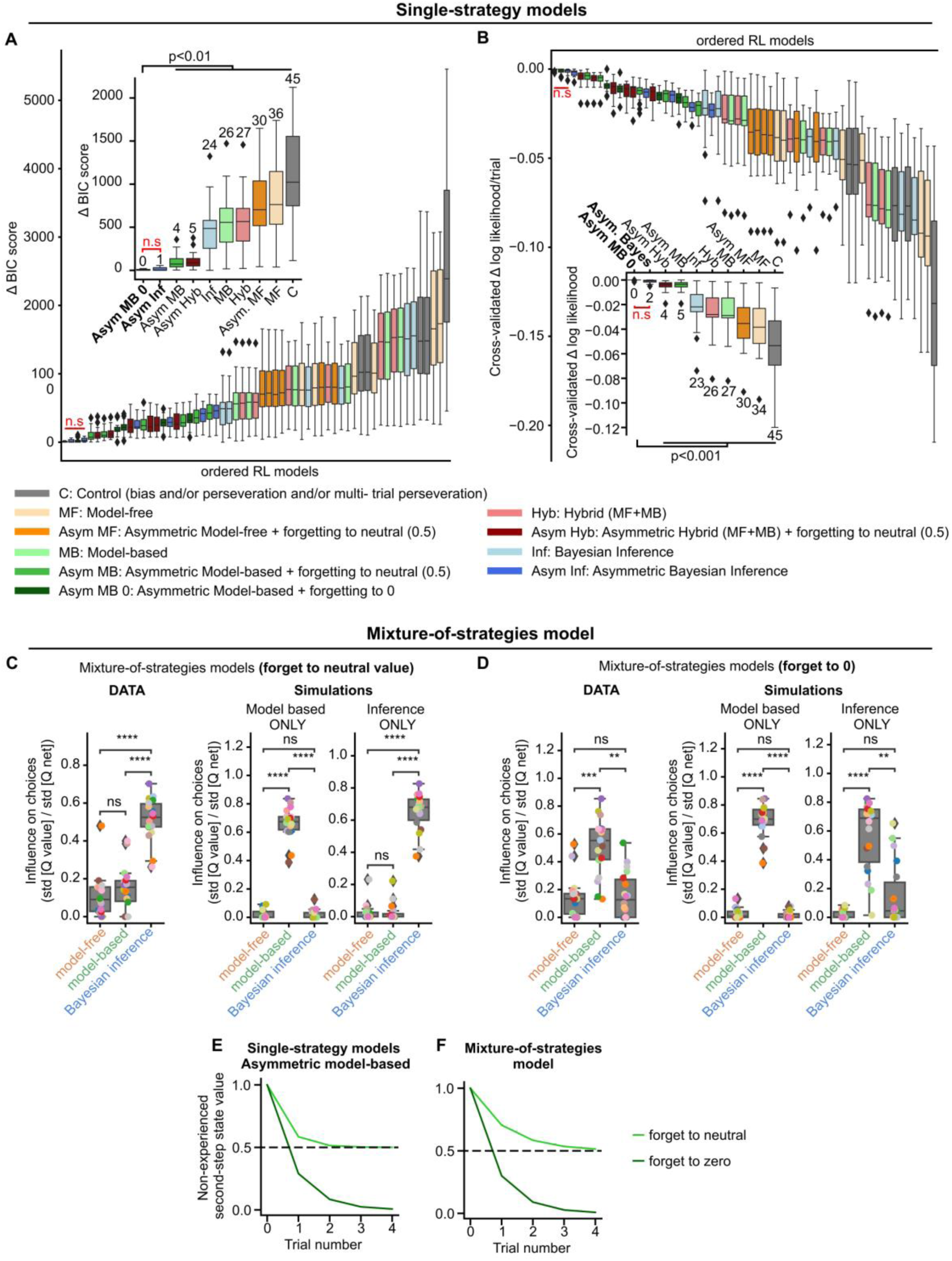
Model comparison and mixture-of-strategies model. **A-B)** Model-comparison for single-strategy models **A)** BIC score model-comparison of all the models tested (see Supplementary Table 2 for details of each model). Colours indicate the model’s strategy. Each strategy was tested both with and without choice bias and multi-trial perseveration. Additionally, models were tested with both symmetric and asymmetric learning from reward and omission outcomes (see Methods). Model-free and model-based RL strategies were tested with and without forgetting about the values of not-visited states and not-chosen actions. Two variants of forgetting were tested: one implemented forgetting as value decay towards a neutral value of 0.5, the other as value decay towards 0. Insets show the best-fitting model of each model class. Fitted parameter values for the best model in each class are presented in Supplementary Table 1. **B)** Cross-validated log likelihood of the same models as in **A**. Inset, as in **A**, best fitting model of each category. ***Both model comparison approaches indicate that asymmetric inference and asymmetric model-based RL with forgetting to 0 strategies explain the data equally well, and better than other strategies considered.* C-F)** Mixture-of-strategies model: choices were determined by a weighted combination of the asymmetric versions of model-free RL, model-based RL, and Bayesian inference. We tested two different mixture-of-strategies models differentiated by how forgetting was implemented for the RL components: In **C, E)** forgetting was implemented as value to decay towards a neutral value of 0.5, while in **D, F)** forgetting was implemented as value decay towards 0. **C)** Mixture-of-strategies model with forgetting towards neutral value. Left panel: influence of each model component on choices, evaluated using the fit of the model to each subject’s data. The influence on choices was quantified as the standard deviation across trials of the difference between the two first-step action values due to a given component, divided by the total standard deviation of the value difference due to all components. Each coloured dot represents a single mouse. Centre and Right panels: influence of each model component for fits of the mixture-of-strategies model to behaviour simulated from either a model-based (centre panel) or Bayesian inference (right panel) single-strategy model with parameters fit to each subject’s choices. The fit correctly assigned high weight to the strategy that generated the simulated data and low weight to the other strategies. **D)** As in **C** but with forgetting towards 0. When forgetting was implemented as decay towards 0, the mixture-of-strategies model fit to simulated data did not correctly identify which strategy generated simulated data. ***Therefore, consistent with the model comparison, we cannot arbitrate between asymmetric inference and asymmetric model-based RL with forgetting to 0 strategies using the mixture-of-strategies model.* E, F)** Value decay of the non-experienced second-step state over consecutive trials when forgetting decayed towards neutral (light green) or zero (dark green) in **E)** the asymmetric model-based model and **F)** in the mixture-of-strategies model. Boxplots show the distribution of the data across subjects. Grey box represents the interquartile range, with horizontal lines representing first quartile, median and third quartile, from bottom to top. Whiskers represent minimum and maximum values. Rhomboids mark outliers. n.s. non-significant, *p<0.05, **p<0.01, ***p<0.001, paired t-test and Bonferroni correction.

**Supplementary Figure 2.**
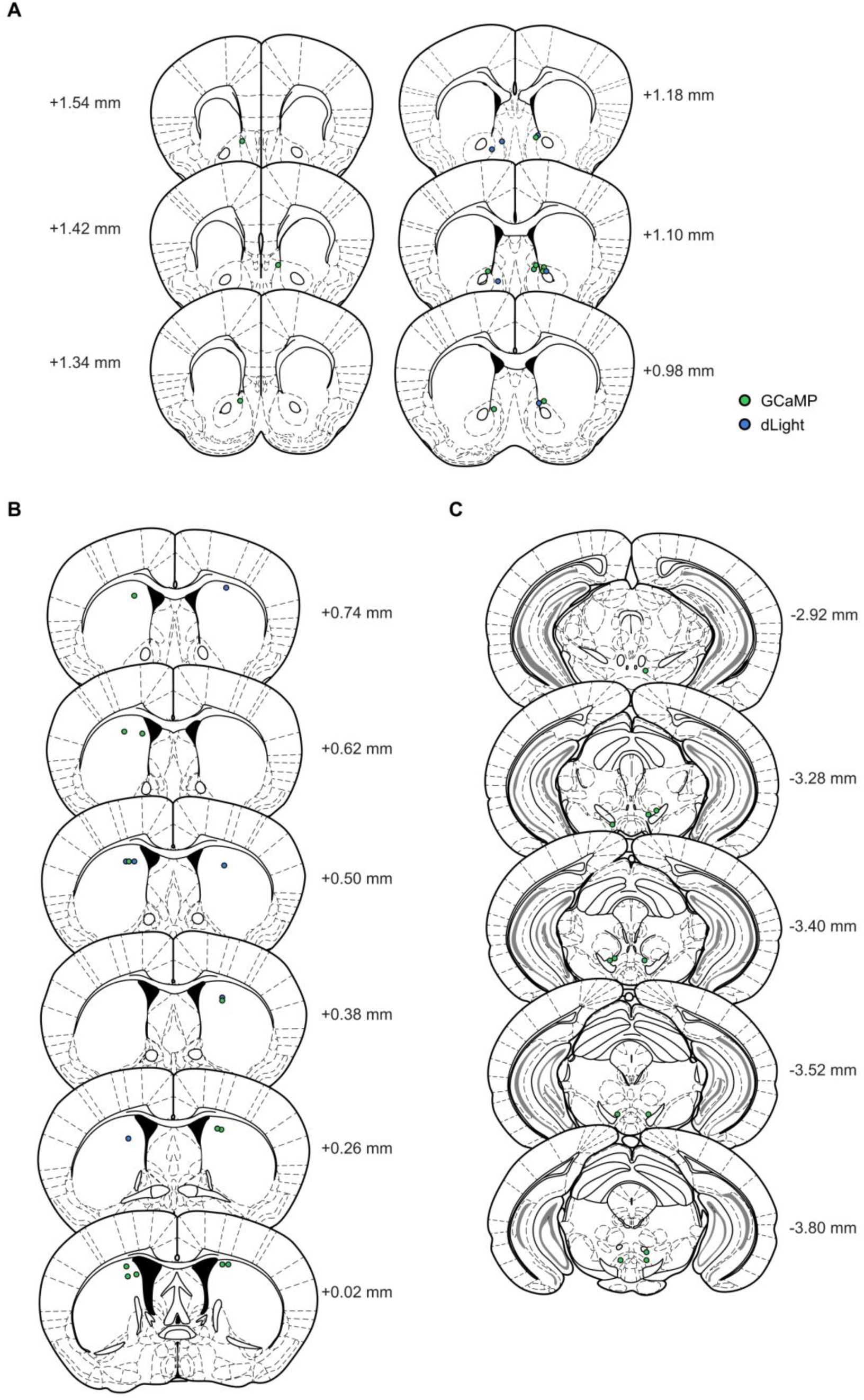
Fibre placement for photometry experiments. **A) NAc,** B) **DMS, and C)** VTA. Green, GCaMP6f; blue, dLight1.1 animals.

**Supplementary Figure 3.**
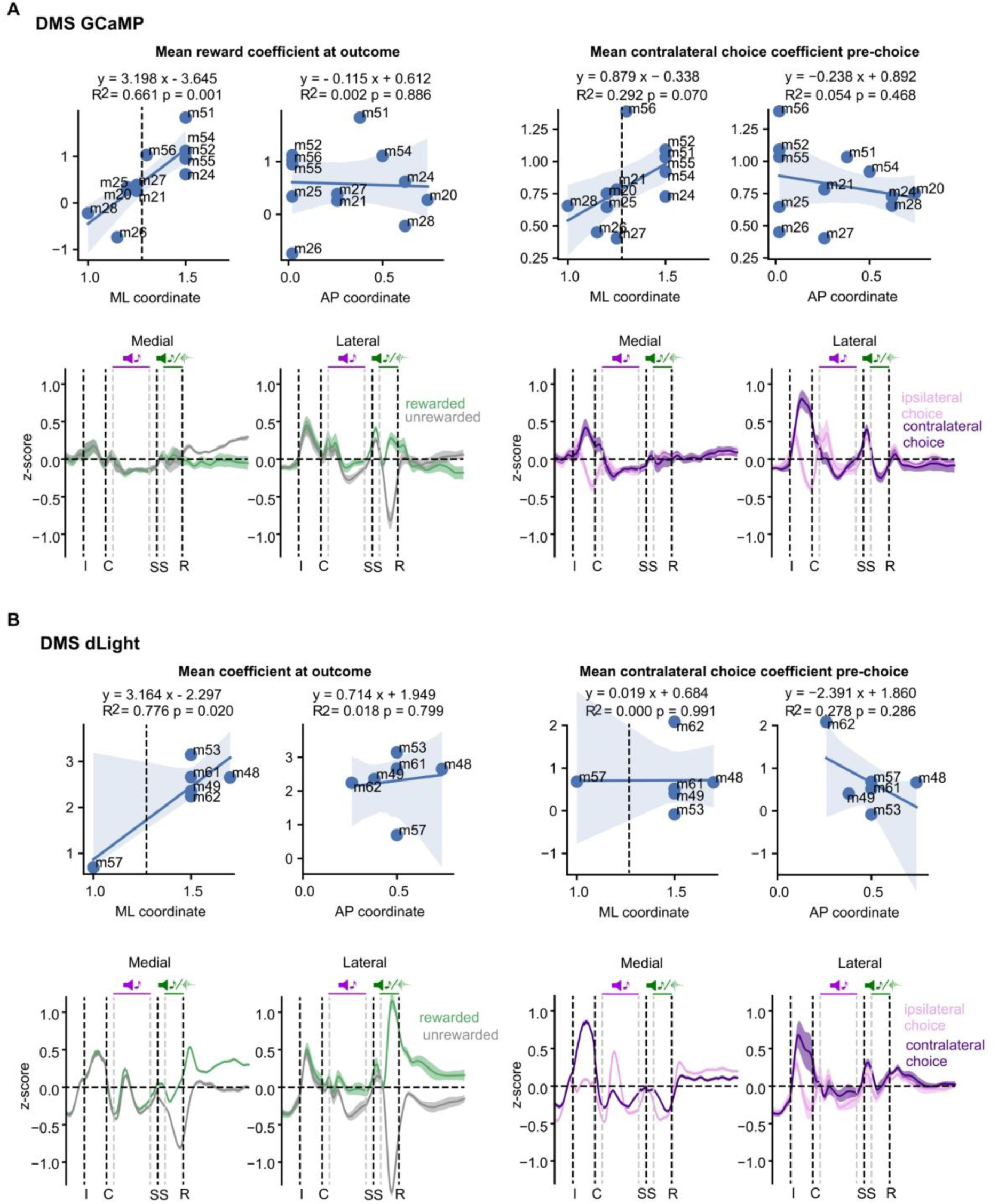
Medio-lateral gradient in reward modulation in DMS. **A)** Dopamine activity measured through GCaMP. **B)** Dopamine receptor binding measured through dLight. *Left*, mean reward coefficient (from the linear regression model) at the time of outcome cue; *Right*, mean contralateral choice coefficient pre-choice. *Top*, correlation between mean coefficient and medio-lateral (ML) or antero-posterior (AP) location of the optic fibre. *Bottom*, mean z-scored activity split by outcome or lateralised choice for animals with medial or lateral placement in DMS. Dashed line in the correlation plots shows how animals were divided into two groups (medial and lateral).

**Supplementary Figure 4.**
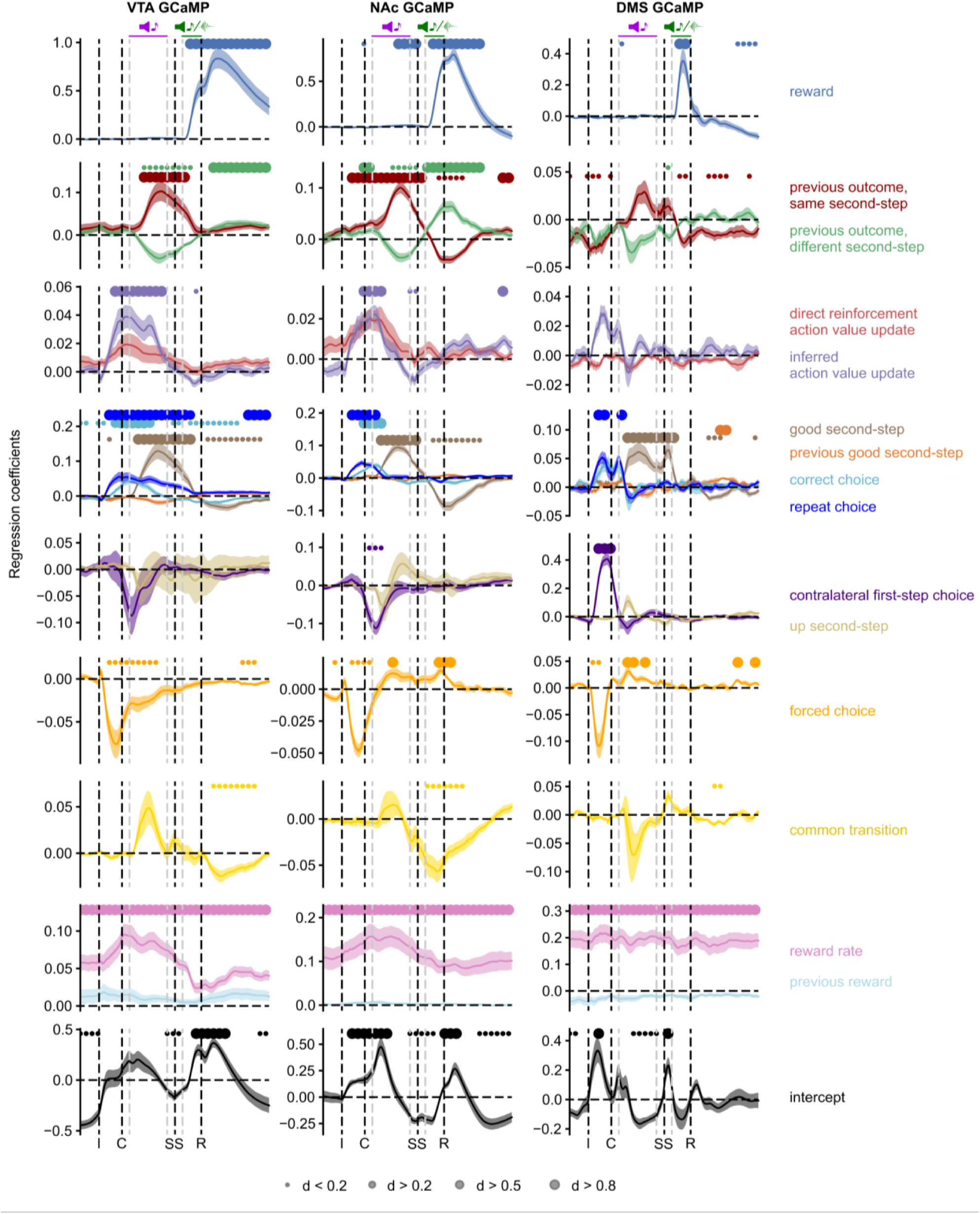
Regression coefficients from all the predictors used in the behavioural logistic regression model predicting VTA, NAc and DMS dopamine activity (GCaMP recordings). Shaded area indicates cross-subject standard error. Shaded area indicates cross-subject standard error. Dots indicate effect size of the statistically significant timepoints, two-sided t-test comparing the cross-subject distribution against 0, after Benjamini-Hochberg multiple comparison correction.

**Supplementary Figure 5.**
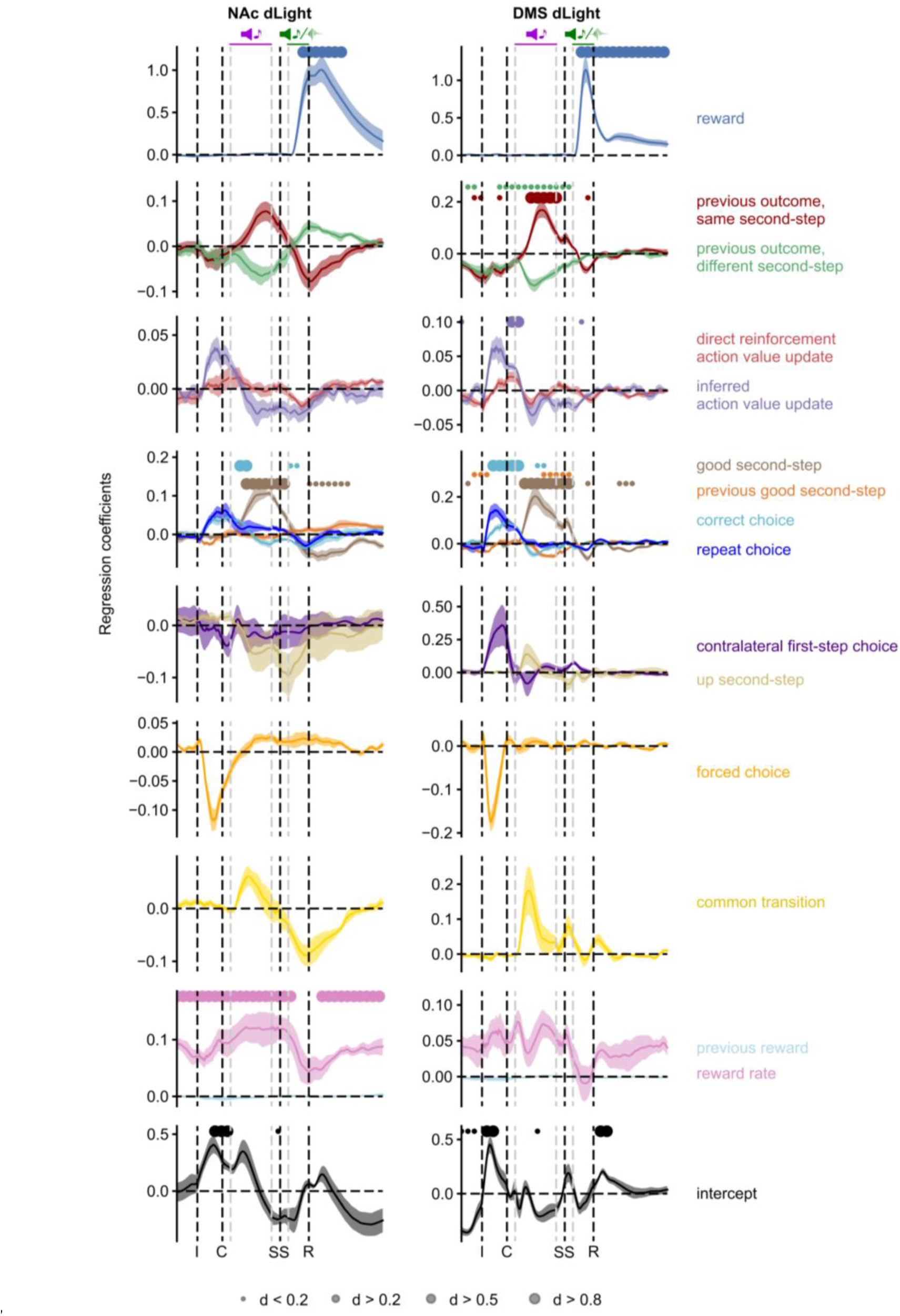
Regression coefficients from all the predictors used in the behavioural logistic regression model predicting NAc and DMS dopamine receptor binding (dLight recordings). Shaded area indicates cross-subject standard error. Dots indicate effect size of the statistically significant timepoints, two-sided t-test comparing the cross-subject distribution against 0, after Benjamini-Hochberg multiple comparison correction.

**Supplementary Figure 6.**
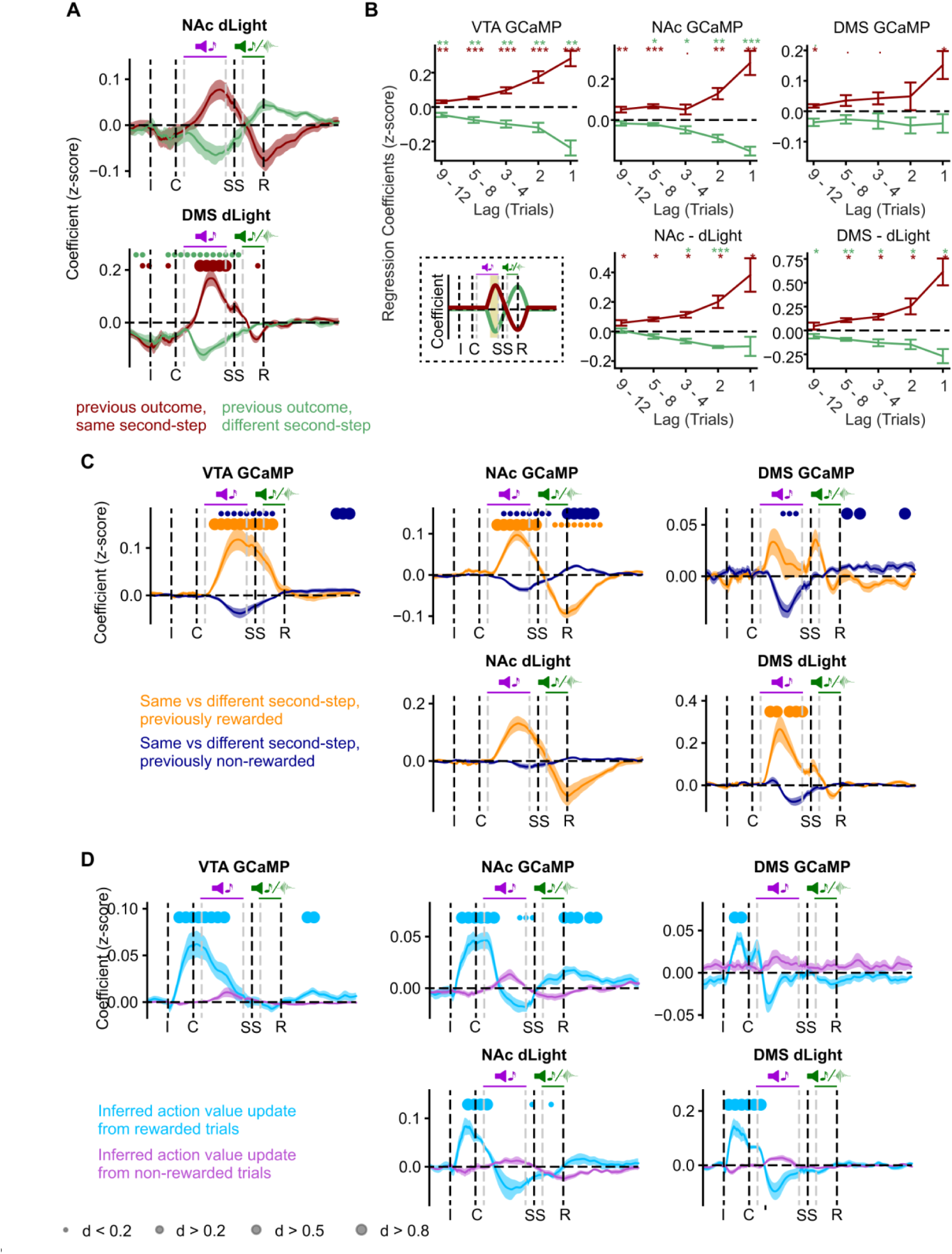
Value update regressors. **A)** Coefficients in the linear regression predicting dopamine activity showing the influence of previous trial outcome when the second-step was the *same* (dark red) or *different* (green) from the previous trial for NAc and DMS dLight signals. **B)** Linear regression predicting the dopamine response to the second-step cue as a function of the extended history of trial events over the previous 12 trials. Bottom left, Schematic of dopamine signal, yellow shaded area represents the predicted activity. **C)** Second-step value update regressors, whether the current second-step was the same or different from the previous trial, split by rewarded and not rewarded outcome on the previous trial. **D)** Model-based action value update split by outcome on the previous trial. Shaded area indicates cross-subject standard error. Dots indicate effect size of the statistically significant timepoints, two-sided t-test comparing the cross-subject distribution against 0, after Benjamini-Hochberg multiple comparison correction.

**Supplementary Figure 7.**
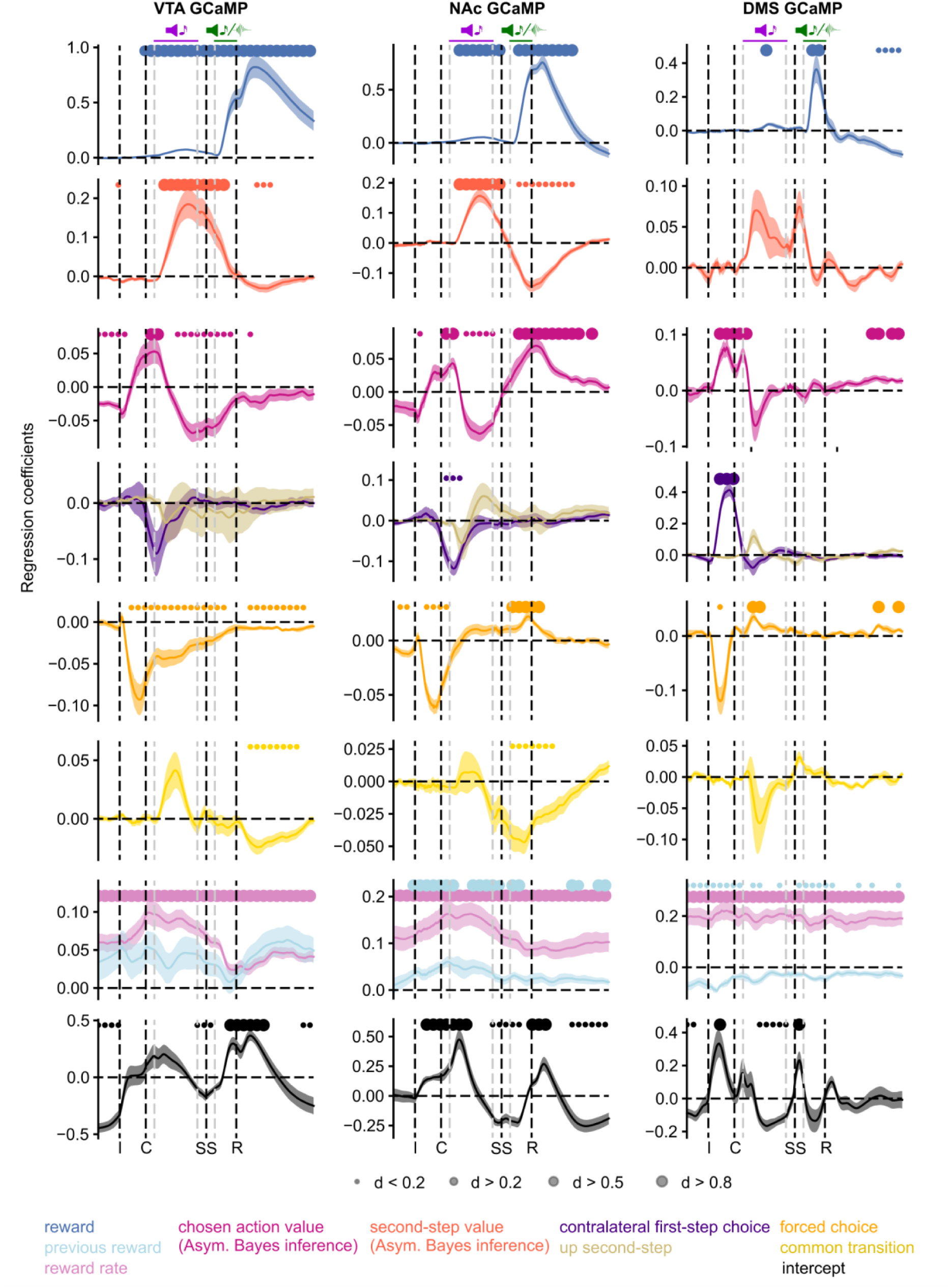
Regression model using second-step and action values derived from the asymmetric Bayesian inference model predicting dopamine activity (GCaMP) in VTA (left), NAc (centre) and DMS (right). Shaded area indicates cross-subject standard error. Dots indicate effect size of the statistically significant timepoints, two-sided t-test comparing the cross-subject distribution against 0, after Benjamini-Hochberg multiple comparison correction.

**Supplementary Figure 8.**
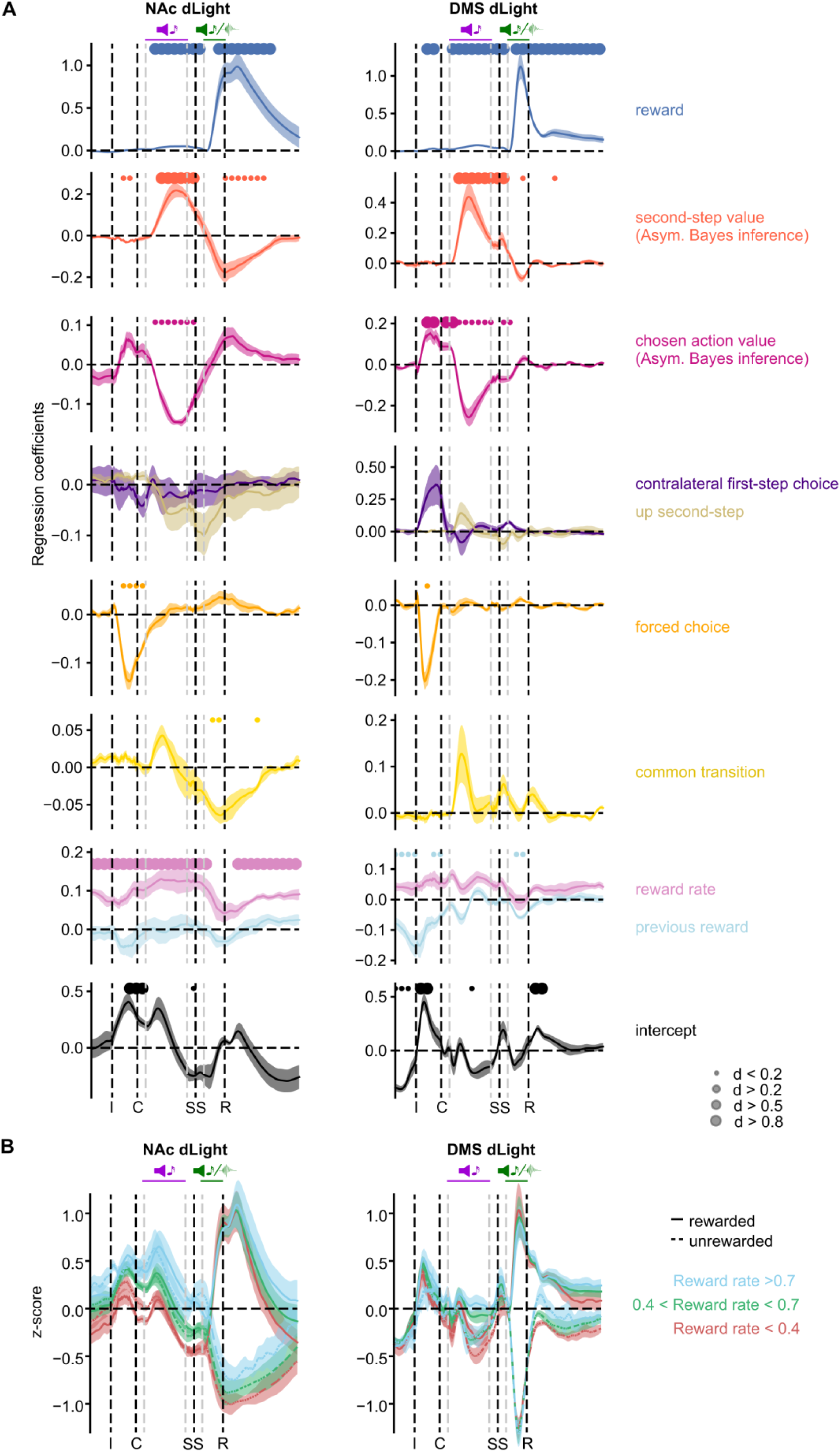
Regression model and reward rate effect in dopamine concentrations. **A)** Regression model using second-step and action values derived from the asymmetric Bayesian inference model predicting dopamine concentrations (dLight) in NAc (left) and DMS (right). Shaded area indicates cross-subject standard error. Dots indicate effect size of the statistically significant timepoints, two-sided t-test comparing the cross-subject distribution against 0, after Benjamini-Hochberg multiple comparison correction. **B)** Mean z-score activity split by rewarded/unrewarded trials and recent reward rate. Blue – high reward rate, > 0.7 rewards/trial (exponential moving average with tau = 8 trials). Green, medium reward rate (between 0.4 and 0.7). Red, low reward rate (< 0.4).

**Supplementary Figure 9.**
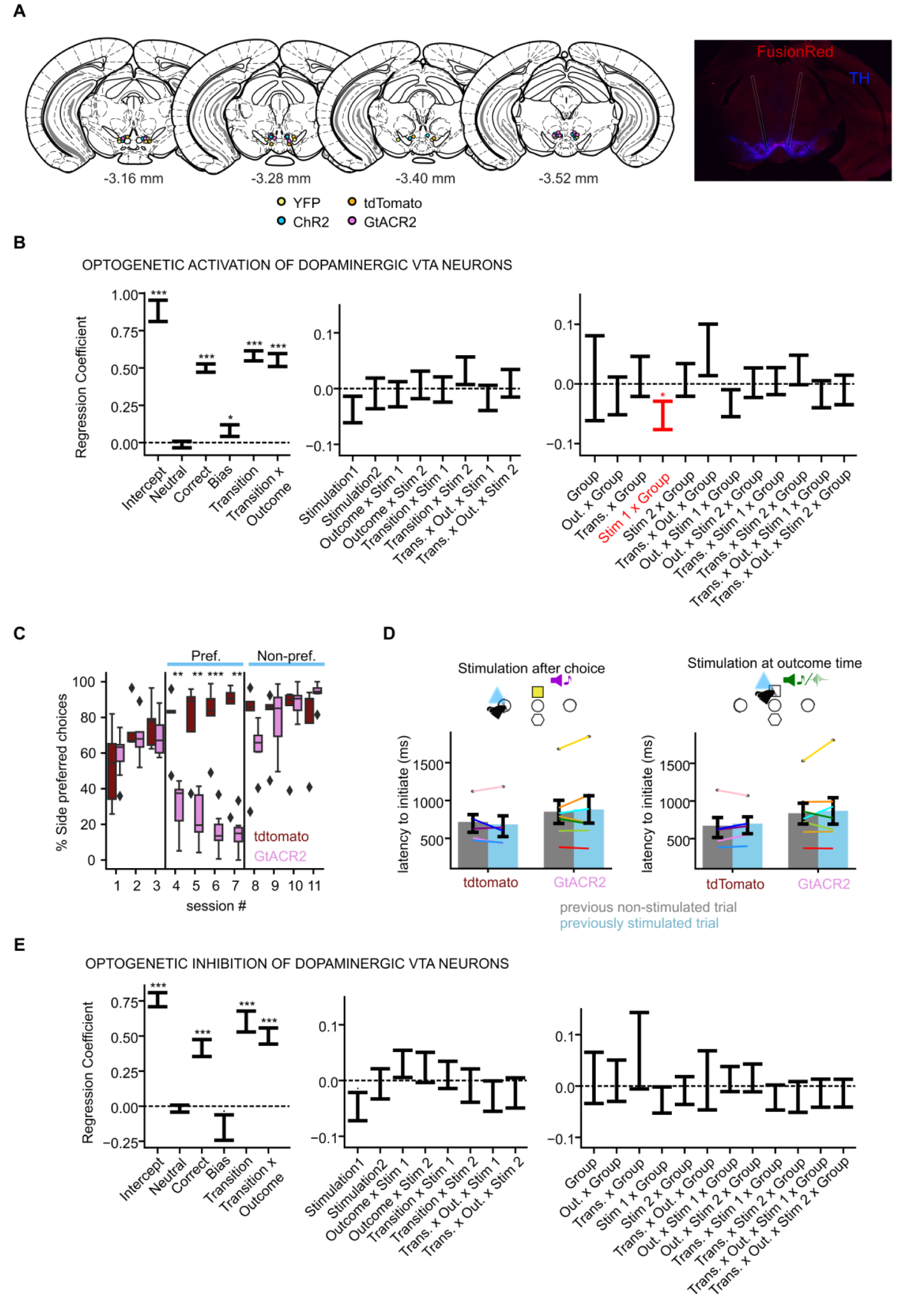
Optogenetic manipulation. **A)** Left, fibre placement. Yellow, YFP animals from the optogenetic activation experiment; blue, ChR2 animals; orange, tdTomato animals from the optogenetic inhibition experiment; pink, GtACR2 animals. Right, photomicrograph showing injection and fibre placement. Photomicrograph comes from an example GtACR2 mouse, stained for TH (Tyrosine Hydroxylase, blue) and FusionRed (red fluorescent protein, red). **B)** Mixed-effects logistic regression predicting stay/switch behaviour in the optogenetic activation experiment. Analysis included data from both groups (YFP and ChR2) and three stimulation types (non-stimulated, stimulation after first-step choice, and stimulation at outcome). **C-E)** Optogenetic inhibition experiment. **C)** 2-alternative forced choice control task showing percentage of choices to the initially preferred side in the tdTomato (red) and GtACR2 (pink) groups. Following baseline sessions without stimulation (sessions 1-3), optical stimulation (1s continuous, 5mW) was delivered from sessions 4 to 7 when mice poked their initial preferred choice (preferred side in sessions 1-3). From sessions 8 to 11 the side of optical stimulation was reversed, now to be coincident with choice of their initial non-preferred side. **p<0.01, ***p<0.005, two-sided t-test with Bonferroni multiple comparison correction. **D, E)** Inhibition effects on the two-step task. **D)** Mean latency to initiate a new trial after the centre poke illuminates following stimulated and non-stimulated trials. Dots indicate individual animals, error bars show standard error. **E)** As in **B**, but for the optogenetic inhibition experiment, mixed-effects logistic regression including both groups (tdTomato and GtACR2) and three stimulation types (non-stimulated, stimulation after first-step choice, and stimulation at outcome). N: tdTomato: 5 animals, 12,555 trials (choice-time stimulation sessions); 13,373 trials (outcome-time stimulation sessions); GtACR2: 7 animals, 14,789 trials (choice-time stimulation sessions), 15,491 trials (outcome-time stimulation sessions).

**Supplementary Figure 10.**
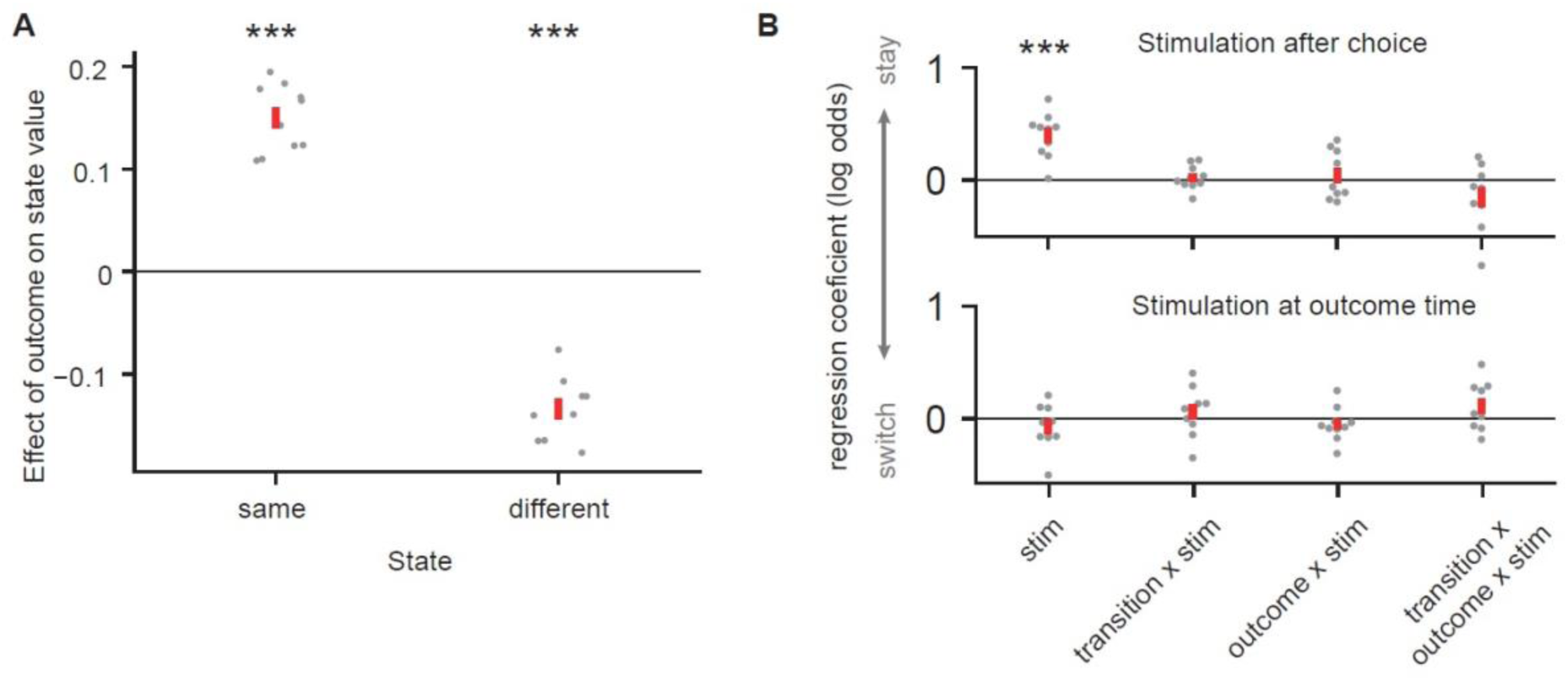
Value updates and simulated dopamine stimulation for version of PFC-basal ganglia network model shown in Figure 6B. **A)** Effect of trial outcome (rewarded vs non-rewarded) on the value of the second-step state where reward was received (same) and on the other state (different). **B)** Effect of stimulated optogenetic stimulation after choice (top panel), or at outcome time (bottom panel). Stimulation was modelled as modifying weights in the basal ganglia network as by a positive RPE.

**Supplementary Figure 11.**
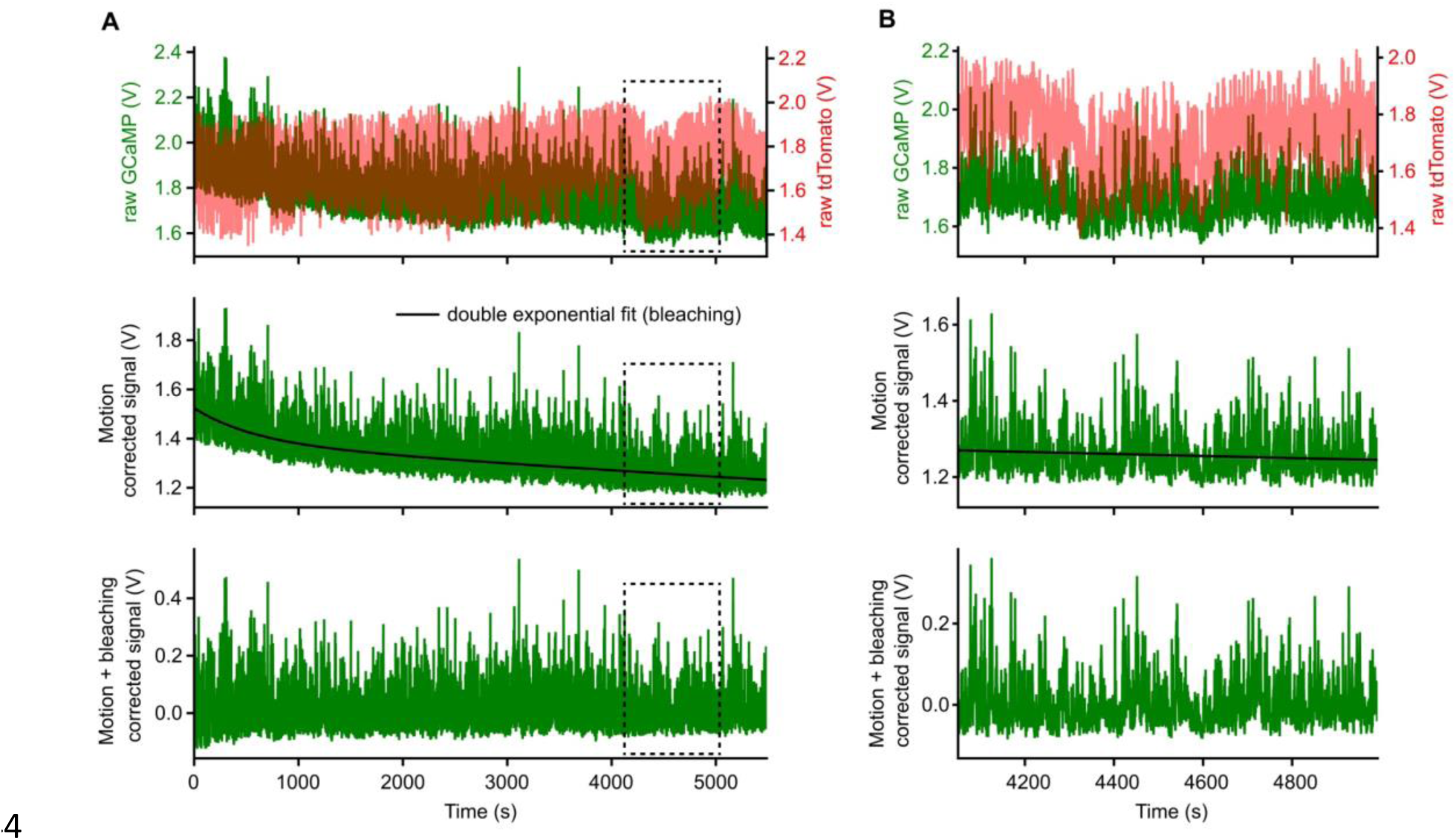
Photometry signal pre-processing. **A-B)** Pre-processing steps in an example session. **A)** Whole recording session. **B)** Zoomed signal from the dashed area in **A**. Top, raw GCaMP and tdTomato signal. Middle, Signal after motion correction, and black line reflects the double exponential fit that will be used for the bleaching correction. Bottom, Motion and bleaching corrected signal.

